# Exploiting variability of single cells to uncover the *in vivo* hierarchy of miRNA targets

**DOI:** 10.1101/035097

**Authors:** Andrzej J. Rzepiela, Arnau Vina-Vilaseca, Jeremie Breda, Souvik Ghosh, Afzal P. Syed, Andreas J. Gruber, William A. Grandy, Katja Eschbach, Christian Beisel, Erik van Nimwegen, Mihaela Zavolan

**Affiliations:** Biozentrum, University of Basel and Swiss Institute of Bioinformatics, Klingelbergstrasse 50–70, 4056 Basel, Switzerland; Department of Biosystems Science and Engineering, ETH Zürich, Mattenstrasse 26, 4058 Basel, Switzerland

## Abstract

MiRNAs are post-transcriptional repressors of gene expression that may additionally reduce the cell-to-cell variability in protein expression, induce correlations between target expression levels and provide a layer through which targets can influence each other’s expression as ‘competing RNAs’ (ceRNAs). Here we combined single cell sequencing of human embryonic kidney cells in which the expression of two distinct miRNAs was induced over a wide range, with mathematical modeling, to estimate Michaelis-Menten (*K_M_*)-type constants for hundreds of evolutionarily conserved miRNA targets. These parameters, which we inferred here for the first time in the context of the entire network of endogenous miRNA targets, vary over ~2 orders of magnitude. They reveal an *in vivo* hierarchy of miRNA targets, defined by the concentration of miRNA-Argonaute complexes at which the targets are most sensitively down-regulated. The data further reveals miRNA-induced correlations in target expression at the single cell level, as well as the response of target noise to the miRNA concentration. The approach is generalizable to other miRNAs and post-transcriptional regulators and provides a deeper understanding of gene expression dynamics.

## Introduction

MiRNAs guide Argonaute proteins to mRNA targets, to repress their expression post-transcriptionally [1]. Transfection of individual miRNAs followed by measurements of transcript and protein levels showed that target mRNA destabilization is an important, possibly the dominant, outcome of miRNA expression [2–4]. However, increasing the time resolution of the experiments, recent studies suggested that translational repression precedes the miRNA-induced target degradation [5–7]. Although a miRNA typically has hundreds of evolutionarily conserved targets [8–10], only very few are down-regulated more than 2-fold in individual experiments [11]. Surprisingly, most miRNA genes are individually dispensable for development and viability, at least in the worm *Caenorhabditis elegans* [12], but the disruption of miRNA biogenesis impairs the ability of embryonic stem cells to differentiate [13]. Thus, strong target repression may not be the primary function of miRNAs [14] and alternative hypotheses need to be explored.

A computational study of small RNA-dependent gene regulation in bacteria initially suggested that post-transcriptional regulators induce thresholds in the response of their targets to transcriptional induction, reducing the effects of spurious transcription [15]. In mammalian systems, miRNA target reporters exhibit a similar behavior [16]. Gene expression being a stochastic process, the number of protein molecules that are produced from a given gene varies between individual cells in a population. Cell-to-cell variability (also called ‘noise’) in protein expression - defined as the variance to mean ratio of the number of protein molecules per cell - depends on the ratio between the rates of mRNA translation and mRNA degradation [17]. Intriguingly, these are precisely the rates that are modulated by miRNAs in a coherent manner, such as to reduce protein expression noise. Consistently, a recent study reported that the variability (coefficient of variation) of the miRNA target CD69, across developing mouse thymocytes is reduced in the presence of miRNAs [18]. The reduction of protein noise is predicted to scale as the square root of the fold-change in protein expression induced by the miRNA [19,20]. This being small for most miRNA targets measured in individual experiments, the relevance of miRNA-dependent reduction of gene expression noise for cellular physiology remains to be established.

As miRNAs can simultaneously bind many mRNA targets, it has been proposed that they provide a ‘channel’ through which target transcripts ‘communicate’ as ‘competing RNAs’ (ceRNAs) [21,22]. Yet for the same reason, it has been difficult to envision how the expression of a single gene, yielding 2-10 transcripts per cell on average [23], can affect the expression of all other genes. Rough estimates of miRNA ‘target abundance’ are in the range of ~10^5^ sites per cell, much higher than the number of cognate miRNA molecules [24]. In this regime of target excess, further over-expressing a miRNA target could not appreciably affect the expression of other targets. It remains possible that occasionally, a miRNA target may reach a relatively high proportion of the transcriptome to function as ceRNA [21,25–27]. These ‘back-of-the-envelope’ calculations did not consider the possibility that targets may not be equivalent in their ability to bind and sequester miRNAs. A computational analysis suggested that miRNA targets may have asymmetric relationships, a high affinity target being able to sequester miRNAs from a lower affinity target at comparable target concentrations, but not the other way around [22]. *In vitro* measurements indicate that miRNA target sites can have widely different affinities for the miRNA-Argonaute complex [28], an observation that is supported by measurements of Argonaute dwelling times on individual miRNA target sites [29]. To understand the response of the entire network of miRNA targets to miRNA regulation, *in vivo* measurements of miRNA-mRNA interaction constants are necessary.

Taking advantage of a system in which the expression of a single miRNA precursor can be induced over a wide concentration range, we measured the transcriptome-level response of ~600 individual cells with mRNA-seq. We obtained experimental evidence for behaviors that were suggested by computational models or evaluated only with miRNA target reporters, including the ultrasensitive response of miRNA targets to changes in the miRNA concentration as well as the change in miRNA target noise in function of the miRNA level. We further uncovered a hierarchy of miRNA targets, defined by the miRNA concentrations where individual targets exhibit the largest change in response to changes in miRNA level. Using the *in* vivo-measured parameters in a computational model of miRNA-dependent gene regulation we confirm previous suggestions that ceRNAs could correspond to targets with low Michaelis-Menten constants. We additionally find that such targets are not uncommon and tend to be highly expressed. Our study reveals substantial differences in the response of individual miRNA targets to miRNAs and provides an approach to characterizing the dynamics of post-transcriptional regulation of mRNA stability *in vivo*, with high resolution, and transcriptome-wide.

## Results

### A system to study the response of an individual cell’s transcriptome by miRNAs

In a previous study [6] we generated human embryonic kidney (HEK) 293 cell lines in which the expression of a miRNA precursor - hsa-mir-199a - and of the green fluorescent protein (GFP) can be induced simultaneously with doxycyline from a pRTSl episomal vector (see Methods and [6]). Of particular interest to our study is the fact that processing of the hsa-mir-199a precursor gives rise to two mature miRNAs, hsa-mir-199a-5p and hsa-mir-199a-3p (**Figure S1**), that differ in their seed sequences and thereby have largely non-overlapping sets of targets (**Figure 1:a**). One of the cell lines contains solely the inducible hsa-mir-199a-encoding pRTSl vector, and we refer to it as i199. The other cell line has in addition a stably-integrated target of hsa-miR-199a-3p, the 3’ untranslated region (UTR) of kinectin 1 (KTN1) downstream of the renilla luciferase coding region. We refer to this cell line as i199-KTN1. The bidirectional nature of the promoter in the pRTS1 vector was characterized before, with protein-coding constructs and fluorescence-activated cell sorting [30]. We cannot employ the same strategy here because the construct encodes a pre-miRNA instead of a protein. Nevertheless, we quantified the miRNA and GFP mRNA expression in populations of cells that we induced with different concentrations of doxycycline, by quantitative RT-PCR, and found that they are highly correlated (**Figure 1:b**, Spearman r = 0.91, p=1.74E-07). This opens the possibility of using the GFP mRNA as a ‘proxy’ for the miRNAs to study the response of miRNA targets to increasing miRNA concentrations in individual cells.

**Figure 1.**
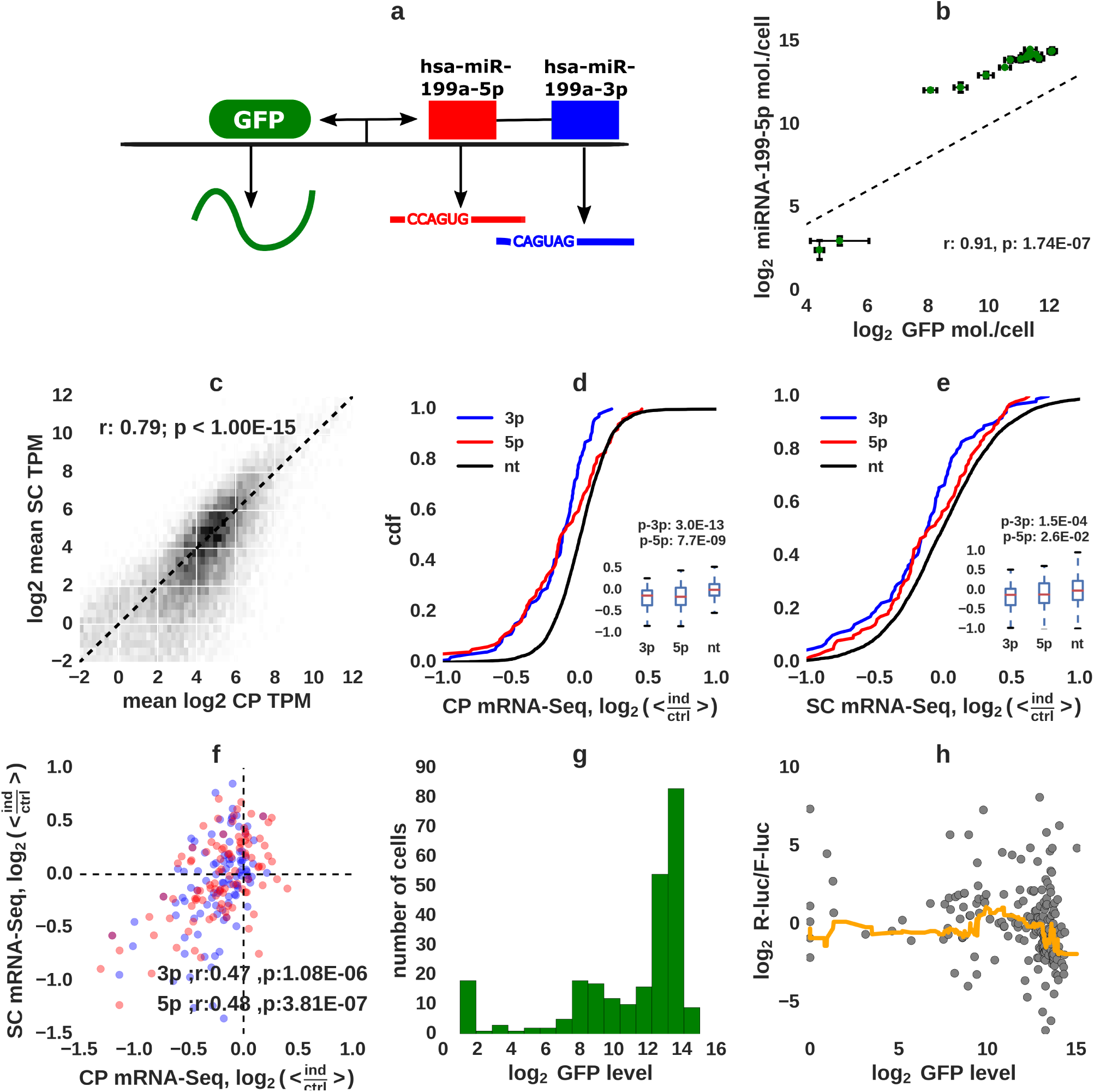
System design and summary of miRNA activity in single cells. **a)** Sketch of the construct used to express hsa-miR-199a-5p (red) and hsa-miR-199a-3p (blue) as well as the reporter GFP mRNA from a bidirectional promoter. Shown are also the ‘seed’ sequences (nucleotides 2-7) of the two miRNAs. **b)** Correlation of hsa-miR-199a-5p and GFP mRNA expression measured in cell populations by quantitative PCR. **c)** Correlation of individual mRNA expression levels (molecules/cell) measured with SC (~40 cells that did not express the miRNAs) and CP (6 replicates of non-induced cell populations) mRNA-seq. Downregulation of top 100 targets of each miRNA as inferred with CP (**d**) and SC mRNA-seq (**e**) (normalized fold changes, blue: targets of hsa-miR-199a-3p, red: targets of hsa-miR-199a-5p, black: other transcripts). **f)** Correlation of expression changes of the top 100 targets of each miRNA in SC and CP mRNA-seq. **g)** GFP mRNA expression distribution (transcripts per million, TPM) in single cells. **h**) Expression (TPM) of renilla luciferase coding sequence with the KTN1 3’UTR, normalized by the expression of the control firefly luciferase. The plot shows the log_2_ ratio in function of GFP expression in each cell (gray dots) and the running median (with window 30, orange line) in function of GFP.

We carried out mRNA sequencing of both single cells and cell populations, and found that the expression levels of individual mRNAs, estimated from cell populations (CP) or by averaging the levels obtained from single cells (SC) that did not express the miRNAs, correlated very well (Spearman r for log_2_ expression values = 0.79, p < 1E-15, **Figure 1:c**). In cells in which expression of hsa-mir-199a was strongly induced for 8 days, CP mRNA-seq revealed the expected down-regulation of MIRZA-G-C-predicted targets [31] (**Figure 1:d**), as did the SC mRNA-seq (**Figure 1:e**). Moreover, we found a moderate correlation between the miRNA target down-regulation inferred with CP and SC mRNA-seq (**Figure 1:f**). To evaluate the response of the entire miRNA target network to miRNA induction, we induced miRNA expression with various concentrations of doxycyline, mixed the induced cell populations and sequenced randomly picked individual cells (**Table S1** shows the induction ranges). The estimated number of GFP transcripts (transcripts per million, TPM, see Methods) obtained from these cells covered a very broad range, from zero to tens of thousands (**Figure 1:g**). Based on the correlation of miRNA and GFP mRNA expression (**Figure 1:b**), we expect a similar range of miRNA levels. We replicated these analyses in the second cell line, i199, with very similar results (**Figure S2**). Because the i199-KTN1 cell line expressed a validated target of hsa-miR-199a-3p, KTN1, we calculated the expression level of this reporter gene in function of miRNA expression and found that KTN1 is down-regulated as the miRNA expression in the cell increases (**Figure 1:h**), as expected.

### Noise and correlation of target expression

The dynamics of small networks composed of miRNAs and targets has been investigated with stochastic models [22,32]. Bosia et al [32] predicted that the coefficient of variation (C_V_) of miRNA targets increases with the transcription rate of the miRNA, showing a local maximum in the region where the miRNA and targets are in equimolar ratio. Similarly, the correlation of expression levels of mRNAs that are targeted by the same miRNA exhibits a maximum around the same threshold. We first reproduced this behavior in a stochastic simulation of a very simple system that consisted of four targets of one miRNA (**Figure 2:a**). The parameters of target expression and interaction with the miRNA were chosen such that (1) two targets (a and b) were down-regulated at lower and two (c and d) at high miRNA concentrations, (2) targets spanned a broad expression range and (3) down-regulation was moderate, as generally observed in experiments. We found the expected dependence of mRNA target noise and correlation of target expression on miRNA expression level. Furthermore, individual targets responded most sensitively to the miRNA at different miRNA concentrations, depending on their parameters of interaction with the miRNA (**Figure 2:b,c**).

**Figure 2.**
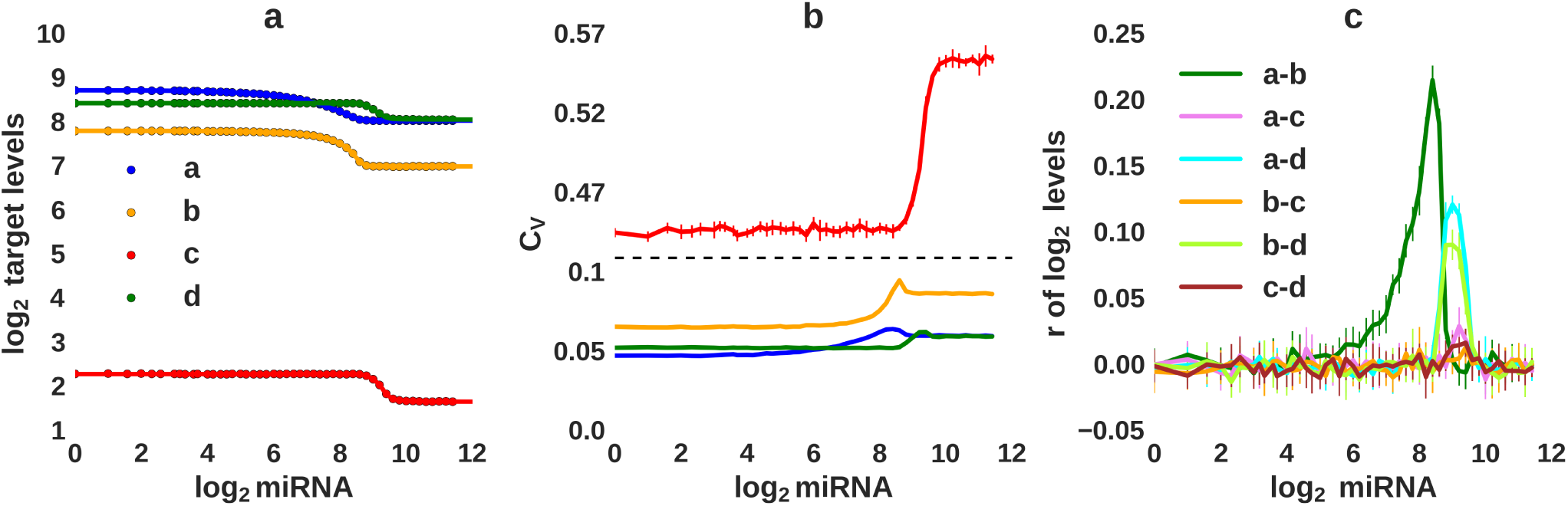
Modeling the effect of a miRNA on multiple targets in single cells. **a)** Results of numerical integration (solid lines) and the average of six stochastic simulations (dots) of a model with four target genes (indicating by color) chosen to cover a wide range of mRNA expression levels and to have either high or low sensitivity to the miRNA. Fifty sets of ‘in silico’ cells, each with a defined miRNA concentration were simulated. **b)** Coefficient of variation (C_V_) of the model genes calculated in function of miRNA expression using the simulation trajectories. **c)** Pearson correlation of expression levels of pair of genes in ‘in silico’ cells shown in function of miRNA expression and calculated from the simulation trajectories.

We then asked whether our experimental system also exhibits these behaviors. We computed the total number of miRNA target transcripts per cell (see Methods for the procedure for transcript selection) and found the expected threshold-decrease in the population of miRNA targets as the number of GFP transcripts (and thereby miRNA molecules) increased (**Figure 3:a and S3:a**). Furthermore, in **Figures 3:b,c** and **S3:b,c** we show the running average of the C_V_, and the running average of the correlation coefficient *r* that were calculated from cells that were sorted according to their GFP mRNA expression. Although the mean C_V_ and mean *r* that we computed from the mRNA-seq data exhibit large fluctuations, particularly in the GFP expression range that was poorly covered by our set of single cells (**Figures 3, S3**), both of these quantities showed peaks at the GFP levels where the downregulation of miRNA targets was most abrupt. In addition, in both cell lines the targets of the two distinct miRNAs that were induced in our system showed consistent behaviors, which would not be expected if they were not coupled through the correlated expression of the two miRNAs **(Figure S4)**. Thus, our experimental findings follow directly the theoretical predictions and the simulation results. We therefore concluded that in spite of the limited precision of the single cell mRNA-seq data, our experimental system allows us to investigate the response of individual targets as well as of the entire target network to miRNA induction in single cells.

**Figure 3.**
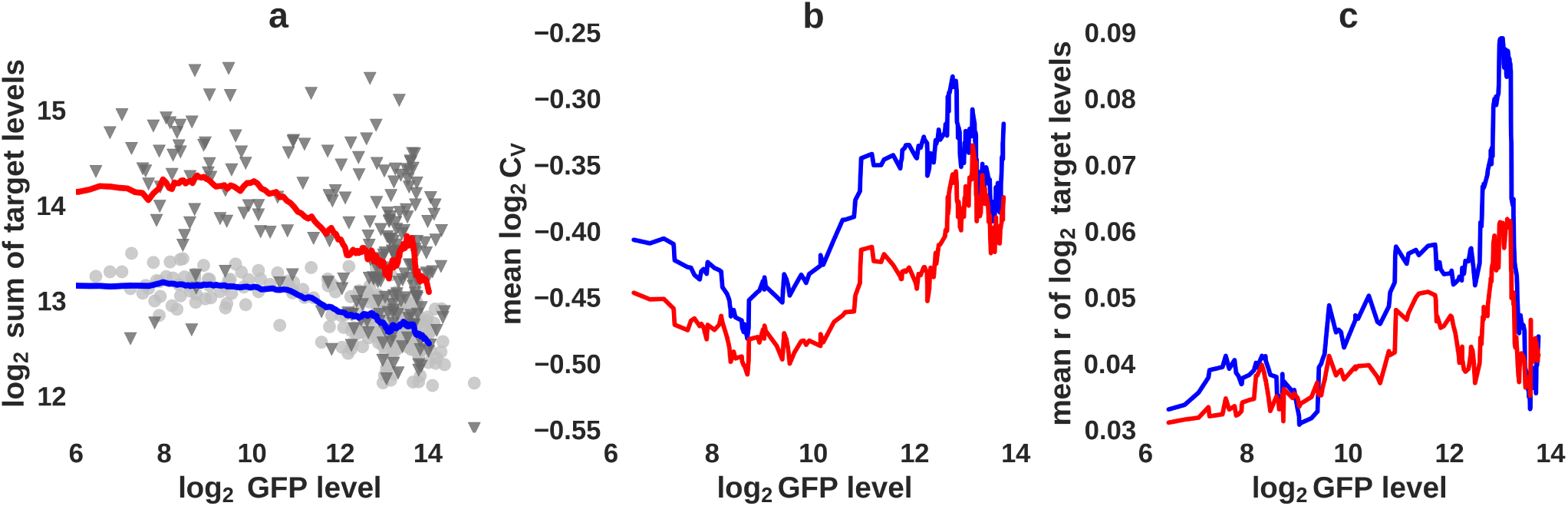
The effect of miRNAs on mRNA targets in single cells. **a)** Total target expression (log_2_ sum of TPMs of hsa-miR-199a-3p and hsa-miR-199a-5p predicted targets, see Methods for target selection) in i199-KTN1 cells in function of log_2_ GFP expression. The colored lines show running means with a window of 30 cells, the grey dots and triangles show the expression of the 3p and 5p targets, respectively, in individual cells. **b)** Mean C_V_ and **c)** mean Pearson pair-wise correlation coefficients for miRNA targets in function of GFP expression. For each GFP level averages were calculated from the fifty cells with GFP expression closest to the reference level.

### Inference of Michaelis-Menten constants of individual miRNA targets

We next asked whether individual mRNAs differ in their response to miRNA induction. As shown in the Methods section (Eq. 1-13) we could infer the Michaelis-Menten constants *K_M_*, that describe the sensitivity of miRNA targets to the miRNA simply from the expression levels of all targets in cells with varying miRNA concentrations. By analogy with enzymatic reactions (see also [28]) *K_M_*s are defined as 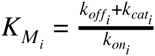, where 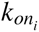 is the rate of association of target *i* with the miRNA-containing effector complex, 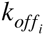 is the rate of dissociation and 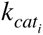 is the rate with which target *i* is degraded when it is complexed with the miRNA (see also [28]). Because the single cell mRNA-sequencing has a fairly high measurement error (see **Figure S9** and [33]), we used simulated data to derive a procedure to smoothen the noisy target levels from single cells and generated the matrix 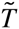 that serves as input for the analytical method. We generated *in silico* data with the model described in the Methods, we added noise to target expression levels such as to match the coefficient of variation corresponding to the technical and biological noise observed in single cell mRNA-seq (**Figure 4:d**) and then we fitted the data to retrieve the input parameters (**Figure 4** and **S7**, and In Silico Model Verification Section). We optimized the fitting procedure by appropriately selecting the cells in which the miRNA expression level was most informative for the change in target levels and by smoothing the *in silico* data to reduce stochastic variations. The final procedure allowed us to robustly and accurately infer the input parameters from *in silico* data (**Figure 4:g,h**). We then applied it to the data from the i199-KTN1 and i199 cells (**Figure 5** and **S5**).

**Figure 4.**
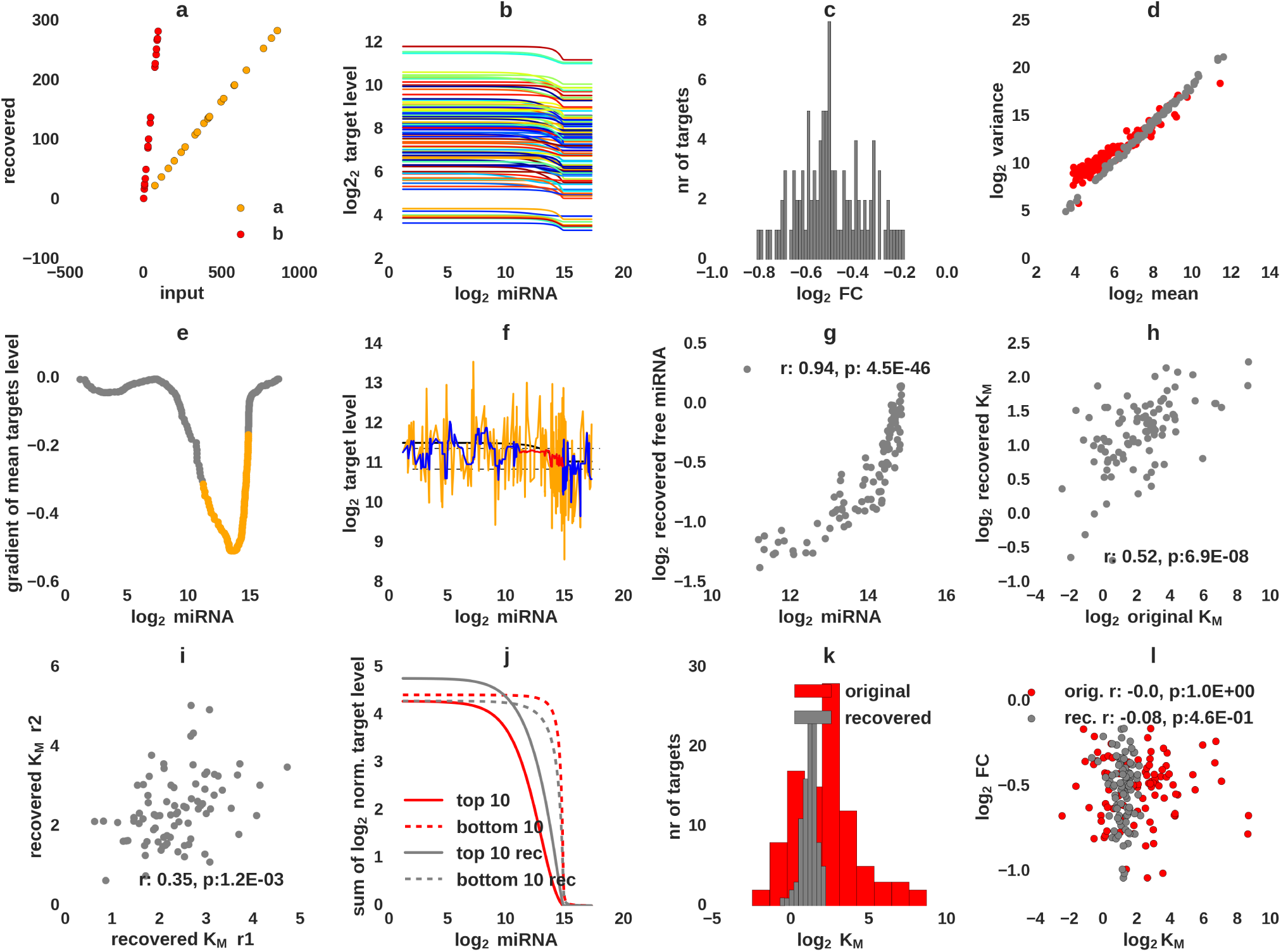
*In silico* data and method verification. **a)** Verification of the reverse Kronecker procedure: We applied the reverse Kronecker procedure for two vectors, **a** and **b**, with random integers. First, we calculated a matrix of ‘expression levels’ as the outer product of the two vectors and the we calculated the solution following Equation 13. The correlation of the input and the inferred values is shown. The values are recovered up to a multiplicative factor. **b)** Generation of *in silico* targets: Expression of the 100 *in silico* targets in function of the miRNA level. **c)** Distribution of log fold-changes 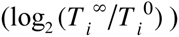 of *in silico* targets at saturating miRNA concentrations. **d)** The log_2_ variance of individual model targets (in grey) shown as a function of their log_2_ mean in individual cells after addition of log-normal noise. For comparison, we show a similar representation for the targets of hsa-miR-199a-3p in KTN1 cells (red). **e)** Selection of *in silico* cells for the inference of *K_M_*s: Cells with a miRNA expression around the point where the total target level changed most in function of miRNA expression (shown with orange) were used to construct the 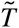 matrix. **f)** Noise filtering: level of an arbitrary target (black) in *in silico* cells with different miRNA concentrations after addition of noise is shown with an orange line. Initially a window of length five was used to calculate a running median (blue) and then an iterative procedure for computing running means was used to ensure that all expression values for gene *i* in cell *j* (the considered cells are shown with orange in panel **e**), T_ij_, are within the interval 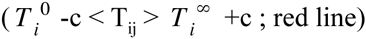, which correspond to the maximum and minimum expression levels of gene *i*. These are achieved when the miRNA is absent and at saturating concentrations, respectively; c is a small fraction of 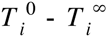 that is used to stabilize reverse Kronecker operation (dashed lines). **g)** Scatter plot of the inferred free miRNA levels and the miRNA levels that were used as input to the model. Spearman correlation value and its associated p-value are also shown. **h)** Scatter plot of the inferred and input log_2_ *K_M_* values. Pearson correlation and its associated p-value are also shown. **i)** Scatter plot of the *K_M_* values that were inferred from *in silico* data that was generated with the same input *K_M_*’s but with two distinct applications of noise to the target expression levels. Pearson correlation and associated p-value are also shown. **j)** Sum of normalized (by corresponding 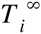 values) log_2_ target levels for ten targets with the lowest (solid line) and highest (dashed line) *K_M_*s. In red, the expression values of targets (without noise) sorted according to model *K_M_*s are shown, while in grey the values of targets sorted according to the recovered *K_M_*s are shown. **k)** Distributions of input (red) and inferred (grey) *K_M_* values. **l)** Scatter plot of *K_M_* values and target fold-changes for input *K_M_* (red) and the inferred *K_M_*(grey) values.

In both cell lines we studied the response of the 200 highest probability targets predicted by MIRZA-G-C [31] for each of the two distinct miRNAs, that were down-regulated at least 10% at the maximum miRNA concentration (log_2_ 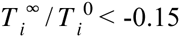, see Methods) (**Figure 5:a**). Based on the gradient of the total target levels in function of GFP we identified the cells where the targets responded most sensitively to the miRNA, and we used them to construct the 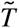 matrix of individual target expression levels in single cells (**Figure 5:b**, see also Equation 10). Next, we selected a few tens of cells with very low GFP and similarly for very high GFP mRNA expression to calculate 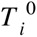 and 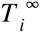, the level of target *i* when the miRNA is not expressed and maximally expressed, respectively. We then inferred the concentration A_F_ of the miRNA in each cell, as well as the Michaelis-Menten constants 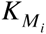 of the interaction between the miRNA and each target *i*. We applied the inference independently to the two miRNAs. We found that the inferred concentration of both free 5p and 3p miRNAs per cell correlated strongly with the GFP level (Spearman r = 0.49, p = 4.8E-8, and 0.65, p = 1.5E-14, respectively, **Figure 5:c**), as expected from the mathematical model and the simulation. Also strongly correlated were the abundances of the two miRNAs (Pearson r = 0.68, p < 1E-15 **Figure 5:d**). We carried out the same analysis in the i199 cell line (**Figure S3**) and found a moderate correlation between the Kms inferred for individual targets in the two distinct cell lines (**Figure 5:e**, Pearson r = 0.42 (p = 0.0041) and 0.38 (p = 0.0026) for the 5p and 3p miRNAs, respectively). These correlations are also in the range that we observed for independent stochastic simulations (**Figure 4:i,h**). Considering the imprecision of the mRNA-seq data, we find these results quite remarkable. Altogether they indicate that our inference is as accurate and robust as possible, given the limited accuracy of the mRNA-seq data. The distributions of *K_M_*s span a similar range of values for both miRNAs and cell lines (**Figure 4:f** and **S3:f**). Simulations suggest that the range may be even larger, because the smoothing of the data appears to narrow the range of inferred *K_M_*s compared to those used as input to the simulation (**Figure 4:k**). Finally, we found a small anti-correlation between the log fold-change of targets and their *K_M_*s, suggesting that high *K_M_* targets undergo a stronger decay, possible due to higher *k_cat_* values (see also next section).

**Figure 5.**
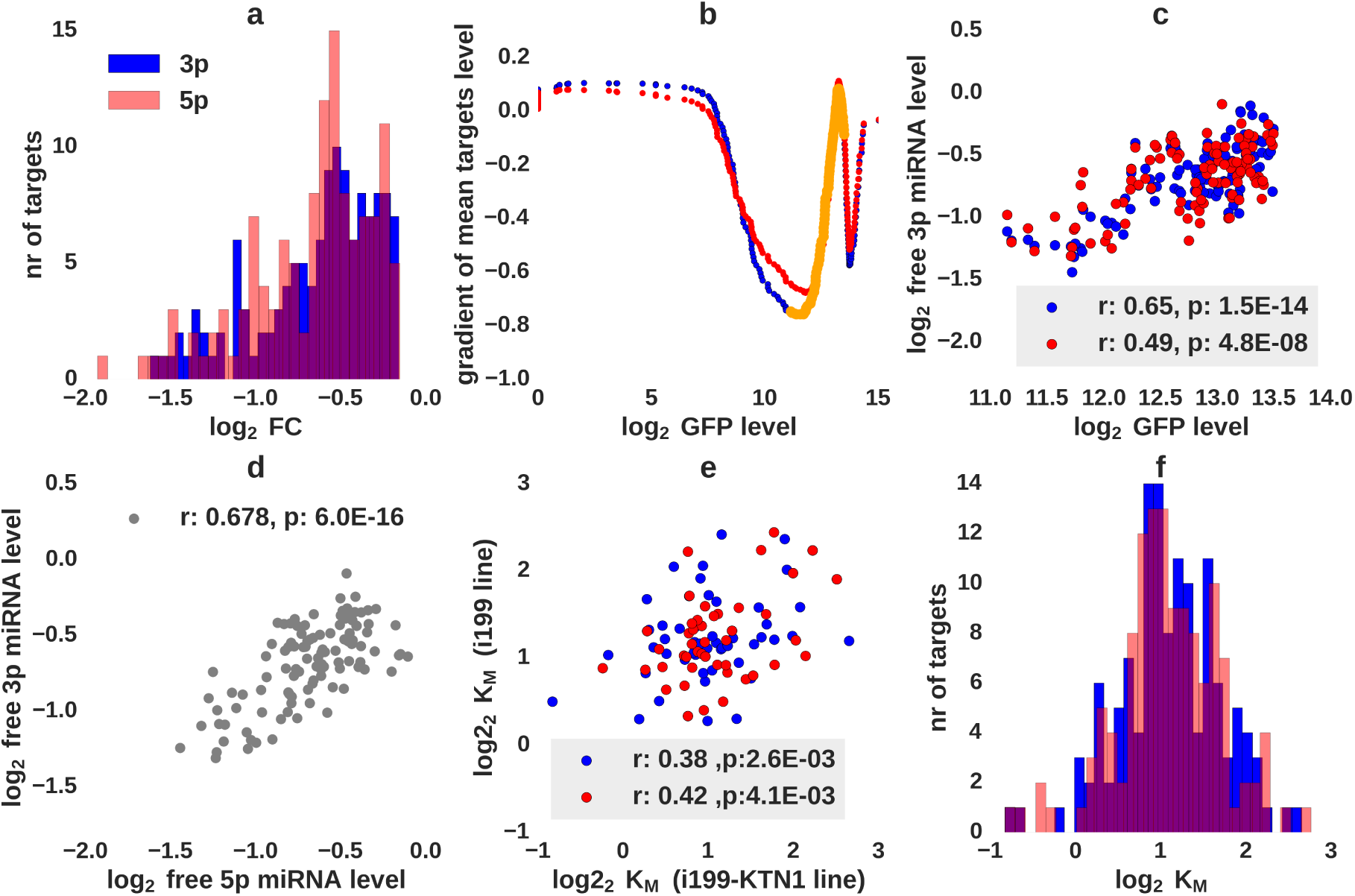
Fitting *K_M_* constants from SC mRNA-seq expression measurements. **a)** Selection of targets. Targets with log_2_ fold change < −0.15 at maximal miRNA concentration were considered. **b)** Selection of cells for the inference of *K_M_*s. Target levels in cells with log_2_ GFP expression in the range of 10-13 (orange line) were used to construct the 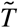 matrix. **c)** Spearman correlation of the inferred levels of free miRNAs 5p and 3p and the measured GFP mRNA. **d)** Pearson correlation of the inferred levels of free 5p and 3p miRNAs. **e)** Pearson correlation of *K_M_* values inferred for individual targets from the i199 and i199-KTN1 cell lines. **f)** Distribution of *K_M_* values inferred for the two miRNAs in i199-KTN1 cells (red: targets of hsa-miR-199a-5p miRNA, blue: targets of hsa-miR-199a-3p miRNA).

### The inferred *K_M_s* define an *in vivo* hierarchy of miRNA targets and shed light on ceRNA function

The behavior of targets with highest and lowest *K_M_*s in function of GFP (and miRNA) expression is shown in **Figure 6:a,b,e** (and **S6:a,b,c**). Targets with low *K_M_*s respond most sensitively at log_2_ levels of GFP of 6-10, whereas targets with high *K_M_*s respond with a steep slope at log_2_ levels of GFP of 12 or more (**Figure 6:a,b** and **S6:a,b**), as expected from the computational predictions (**Figure 4:j**). As an additional validation of the sensitivity of individual targets to the miRNA, we used qPCR to quantify their response to miRNA induction with increasing doses of doxycycline. We selected five targets for which we inferred low *K_M_*s based on the single cell data and five targets for which we inferred high Kms. Of these, two targets did not show a response to miRNA expression in the qPCR assay and one was not detectable. The results for all other seven targets are shown in **Figure 6:c,d**. Although the variability of individual targets is again high, overall, low *K_M_* targets responded at lower miRNA transcriptional induction compared to high *K_M_* targets (**Figure 6:f)**.

**Figure 6.**
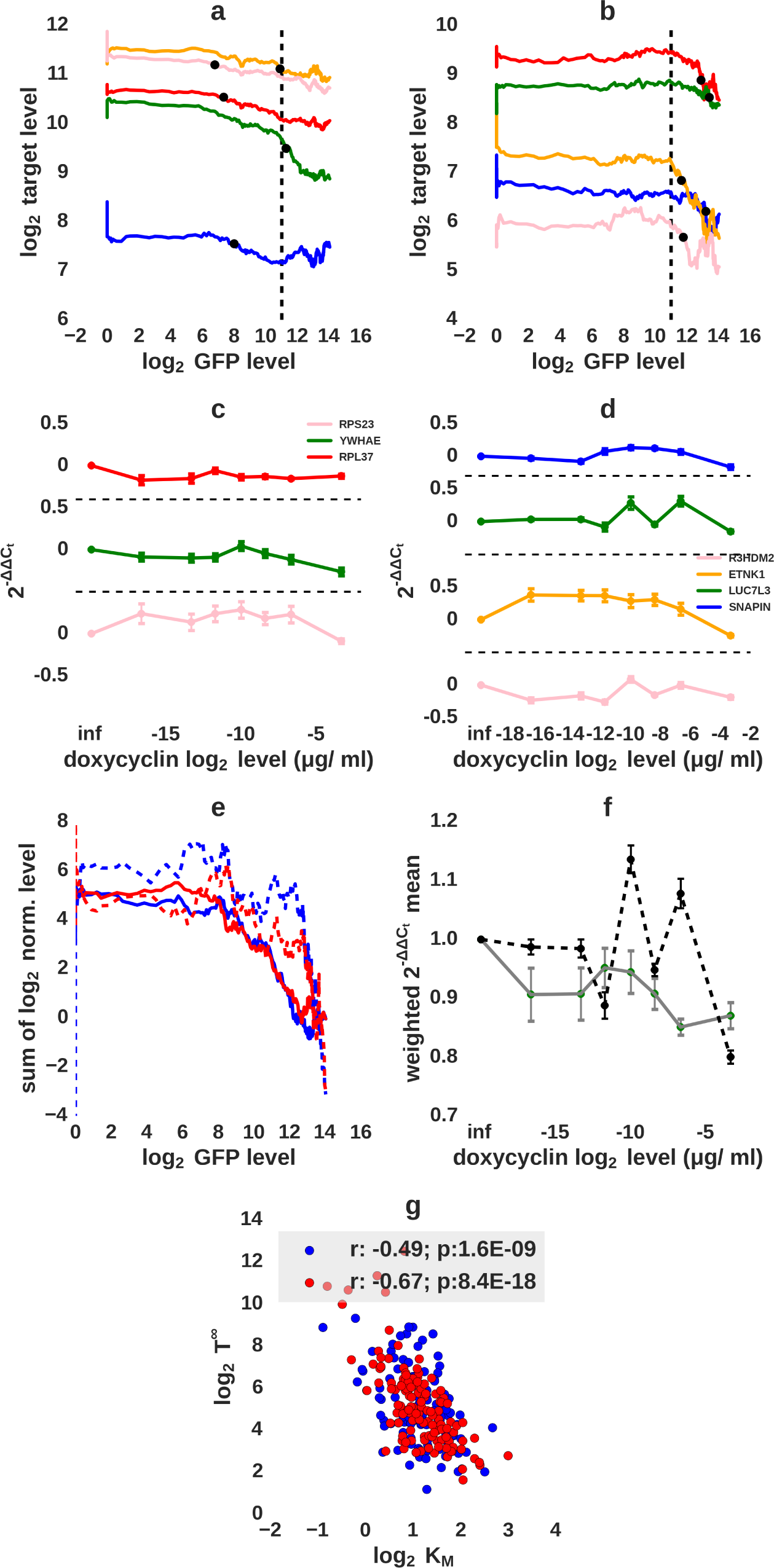
Validation of *K_M_s* fitted based on the single cell data. **a)** Single cell level expression of five low *K_M_* miRNA targets in function of the GFP mRNA. Shown are the running means (window size of 30 cells, black dots show position of the minimum value of the gradient). **b)** As in a, but for five high *K_M_* miRNA targets. **c)** Mean 2^−ΔΔCT^ values (normalized to GAPDH and further to the values in un-induced samples) for three low *K_M_* targets also shown in panel a (colors are preserved). Cells were induced with increasing concentrations of doxycycline and for each concentration, four biological replicates were used to carry out the qPCR. **d)** As in c but for four high *K_M_* targets also shown in panel b (colors are preserved). **e)** Sum of log_2_ running means (normalized before summation by the corresponding 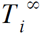 values) for ten targets with the lowest (solid line) and highest (dashed line) inferred *K_M_* values. Red and blue correspond to targets of the hsa-miR-199a-5p and hsa-miR-199a-3p miRNAs, respectively. **f)** Average response of the targets, inferred from qPCR measurements. The weighted means 2^−ΔΔCT^ were calculated separately for low *K_M_* (solid line, targets from panel c), and high *K_M_* targets (dashed line, targets from panel d), using the mean values and standard deviations from biological replicates. Error bars show standard error of the weighted mean (see also methods). **g)** Scatter plot of *K_M_* and 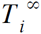 for individual targets of the 5p (red) and 3p (blue) miRNAs. r is the Spearman correlation coefficient.

In the Methods section we derived *K_M_* as 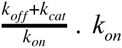 reflects the likelihood of the mRNA meeting a cognate miRNA, which is limited by the diffusion process, and may be similar between targets. If this were the case, the targets with the lowest *K_M_* should be those that have a low dissociation rate *k_off_* (high-affinity), but also a low degradation rate *k_cat_* (see also equation 8), whereas a high *K_M_* could be achieved either with a high dissociation rate or a high rate of miRNA-induced degradation. Thus, we expect *K_M_* to be correlated with *k_cat_* and, because 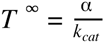, we may expect *K_M_* to be anti-correlated with *T*^∞^, which is indeed what we observe, consistently for the targets of the two miRNAs (**Figure 6:g and S6:d**).

To evaluate the implications of the inferred *K_M_*s for the debate on competing endogenous RNAs [24,34], we used again our computational model with realistic *K_M_* values and explored the effect that one miRNA target (the ceRNA) can have on the expression of all other targets. The parameters of the targets were set as described in section “In silico model verification”. We set the decay rate of the free ceRNA to 0.1/h, and to *k_cat_* = 0.11/h, when complexed with the miRNA. The *k_on_* and *k_off_* parameters of the ceRNA were adjusted to achieve three different values of *K_M_:* 0.05, 1.0 and 25.0, corresponding to a low, intermediate and high *K_M_*. We then asked what the behavior of targets with low *K_M_* (*K_M_*<2M) and high *K_M_* (*K_M_*>2M) is, as the ceRNA is induced transcriptionally. As shown in **Figure 7**, we found that moderate/high transcriptional induction of the low *K_M_* ceRNA can lead to the upregulation of low *K_M_* targets at moderate miRNA concentrations and of high *K_M_* targets at relatively high miRNA concentrations. On the other hand a high *K_M_* ceRNA can only influence the expression of other high *K_M_* targets and only when the cognate miRNA is in high concentration. What has been perhaps under-appreciated so far is that the de-repression the ceRNA can induce is bounded by the repression that these targets underwent in the first place, in the absence of the ceRNA. That is, only targets whose decay is highly accelerated by the miRNA can be strongly de-repressed by a ceRNA.

**Figure 7.**
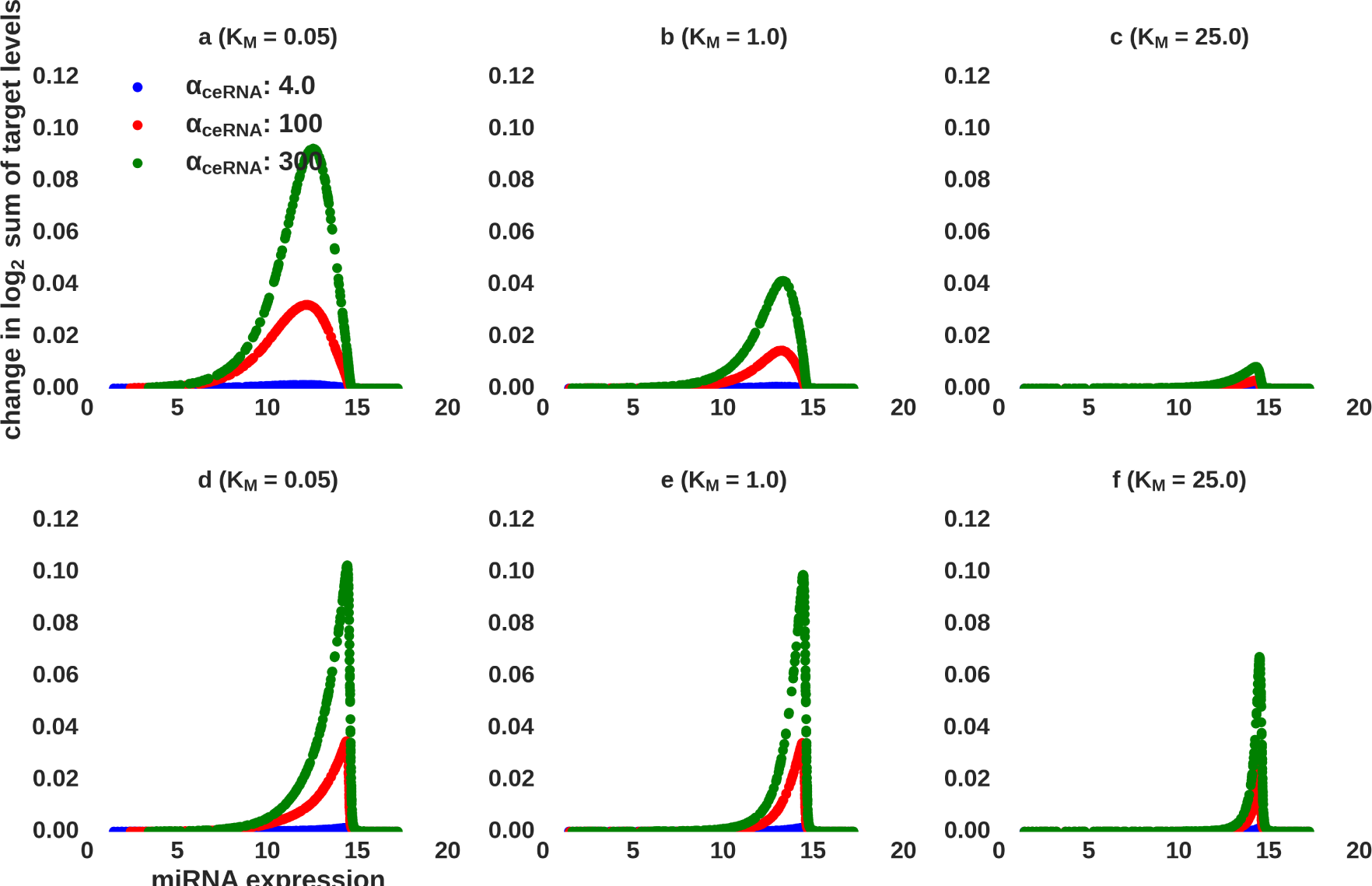
Prediction of the response of different types of miRNA targets to the induction of expression of a ceRNA. A competing RNA with low (**a, d**), medium (**b, e**) or high (**c, f**) *K_M_* is transcriptionally induced at three different levels. **a),b),c)** show the response of targets with *K_M_*<2M to the transcriptional induction of the ceRNA whereas **d),e),f)** show the response of targets that have *K_M_* > 2M. 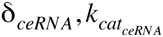 were set to 0.1/h and 0.11/h, respectively.

## Discussion

Single cell RNA-sequencing has opened a new route to understanding how cells regulate themselves and contribute to higher order structures. This technology has been used to characterize transcript isoforms and gene expression [35], to improve classification of cell types [36], and to discover new, particularly rare types of cells [37]. Single cell analysis has also been used to infer parameters of gene expression (see [38] for a recent review), but this methodology has not been applied so far to study the regulation of mRNA stability by miRNAs and its consequences for cellular behavior.

The relatively low capture rate of individual mRNAs and the large technical noise of single cell sequencing remain important issues, but new developments such as mRNA barcodes [39] continue to push the boundary towards ever increasing accuracy. Although miRNAs are viewed as a means to buffer stochastic fluctuations [40], few studies have investigated how miRNA targets respond to miRNA expression at the individual cell level. In particular, some studies estimated the effect of endogenous miRNAs on the protein expression noise of reporter targets with multiple miRNA-complementary sites [16,20]. The reduction in protein expression noise has been related to the degree of miRNA-induced down-regulation, which is generally limited, except for reporters that carry multiple perfectly-complementary miRNA binding sites in their 3’ UTRs. Whether miRNAs regulate the expression noise of natural targets in physiological conditions remains to be determined (see also [20]). Reporter targets have also been used to investigate the cross-correlation and synchronization of miRNA targets in the presence of the cognate miRNAs [41]. The response of the entire miRNA target network of individual cells, where the mRNAs are expressed at physiological levels, has not been investigated so far.

In this study we developed a robust methodology to characterize, in parallel, the regulatory effects of a miRNA on hundreds of mRNA targets in single cells. We constructed an experimental system in which the expression of a miRNA precursor can be induced with doxycycline, together with the expression of GFP, from a bi-directional promoter. In this system, we have reproduced previously reported effects of the miRNA on the variability of expression of its targets, as well as the increased correlation of target expression in the presence of the miRNA. Furthermore, we developed a methodology for variational fitting of Michaelis-Menten *(K_M_)* constants, that characterize the interaction of the induced miRNAs with mRNA targets. This method takes advantage of the variability in transcriptional activity between individual cells, including at the miRNA-expressing locus. For the first time we have uncovered the hierarchy of targets of a miRNA, defined by the concentration at which they respond to the miRNA induction, in a specific cell type. Interestingly, although many studies reported that targets tend to have low expression in cells in which the miRNA is highly expressed [42–44], we found that miRNA targets cover a very wide expression range. They included even highly abundant, ribosomal protein mRNAs. It could be argued that these mRNAs are not the natural targets of the miRNA, because the miRNA that we have induced is not expressed under physiological conditions in the cell type that we studied. To fully address this possibility one would have to carry out a similar experiment, progressively removing a highly abundant, cell type-specific miRNA, rather than inducing miRNA expression in a cell in which the miRNA is not usually expressed. The miR-122 in liver cells could be a good candidate (see also [24]). However, some abundant, low *K_M_* targets, such as the ribosomal protein 37 (RPL37, 5p target) and tyrosine 3-monooxygenase/tryptophan 5-monooxygenase (YWHAE, 3p target) activation protein (**Figure 6:a**) are also ubiquitously expressed and should be exposed to hsa-mir-199a in a physiological setting.

Interestingly, early analyses of miRNA and target expression already revealed that many miRNA targets are expressed at relatively high level in the tissue in which the miRNA is expressed [43], but this observation has been attributed to a function of miRNAs in optimizing protein output rather than entirely suppressing it. Here we found that the expression level of a target and its *K_M_* are anti-correlated, implying that highly expressed targets are down-regulated at lower concentrations of the miRNA compared to lower-abundance targets. This may relate to the suggestion that miRNAs help reduce the expression of abundant targets to ensure rapid and robust changes of gene expression programs at developmental transitions [43]. Thus, applying our methodology to model systems investigating developmental transitions could substantially improve the understanding of the physiological function of miRNAs. Our analysis also suggests that, because low *K_M_* targets are only sensitive to the expression of similarly low *K_M_* targets, they could impose an affinity threshold for miRNA-dependent regulation, which would otherwise affect a large fraction of the transcriptome.

An avenue that we find compelling to follow in the immediate future is to use the *K_M_* constants along with measurements of mRNA expression changes to understand the dynamics of miRNA targets beyond the down-regulation at saturating miRNA concentrations, which is what current miRNA target prediction tools attempt to model. Although in our study we found that *K_M_* values correlated between related cell lines, it remains to be determined how sensitive these target-specific parameters are to the cellular environment in which the target (and miRNA) are expressed. The biggest challenge in applying our approach on an extensive scale relates to the limited coverage and accuracy of single cell sequencing, but the technologies are improving at a very fast rate. Finally, the approach that we introduced here could be easily extended first to RNA-binding protein regulators of mRNA stability and then to other types of regulators such as transcription factors.

## Acknowledgements

We are grateful to Andrea Riba, Alexander Kanitz, Joao Guimaraes, Andreas R. Gruber and the other members of the Zavolan group for providing input and feedback during the project and for the careful reading of the manuscript. The work was supported by the European Research Council Starting grant 310510-WHYMIR and by the SystemsX.ch systems biology initiative in Switzerland through the RTD grants 51RT-0_145680 (StoNets) and 51RT-0_145728 (NeuroStemX).

## Methods

### Theory

To study the transcriptome-wide effects of miRNA expression we use the following model. Let us consider *M* mRNA targets of a given miRNA, each with a specific transcription rate α*_i_*, decay rate δ*_i_*, and concentration *m_i_*, with *i ∊* {1,… *M*}. Target *i* can bind a miRNA-containing Argonaute (Ago) complex at rate 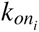, and dissociate from the complex at rate 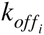. As a first approximation, we neglect the dynamics of the miRNA and assume that the total concentration A of Ago-miRNA complexes in the cell is constant. The concentration of free Ago-miRNA complexes will then be 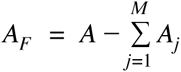. Ago-complexed mRNAs decay at rates 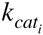. Under this simple model (see also [11]), free mRNAs (*m_i_*) and Ago-bound mRNAs 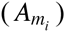 follow the dynamics described by the system of 2*M* differential equations

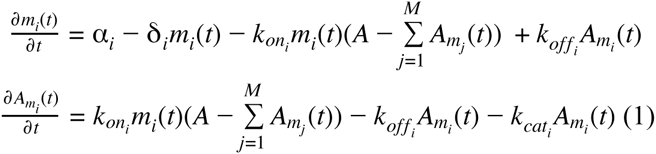

Denoting the total concentration of transcripts *i* (either free or complexed with Ago-miRNA) by *T_i_* and summing the two equations for mRNA *i*, the dynamics of *T_i_* will be described by

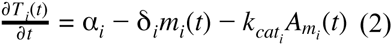

or, in terms of the fraction *f_i_* of molecules of mRNA *i* that are complexed with Ago-miRNAs,

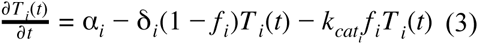

Defining the total concentration of target *i* when no miRNA is present as 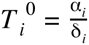 and when Ago-miRNA complexes are in high excess as 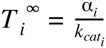 we obtain the total concentration of target *i* at a steady state as

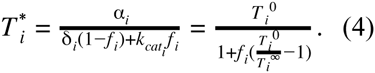

Note that the concentration of the miRNA is reflected in the fraction of bound target. In our experimental system, we vary the expression of the miRNA from very low to very high levels and we can therefore estimate 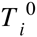 and 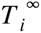. However, the fraction of bound target *i* depends not only on the constants of interaction of target *i* with Ago-miRNA complexes, but also on all other targets that are present in the system. To determine these interaction constants we first derive the *f_i_* s as follows. At equilibrium, we have

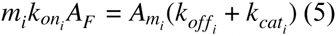

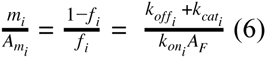

and thus

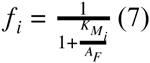

with the Michaelis-Menten parameter defined as 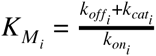. Considering all cells *j ∊* {1,…,*N*}, each with a different concentration of free Ago-miRNA complexes 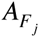, and substituting f in equation (4) we obtain

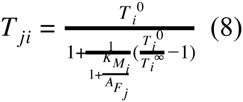

where *T_ji_* is the total concentration of mRNA *i* in cell *j*. We isolate the ratio 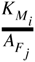 and rewrite

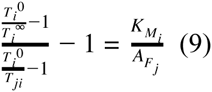

or equivalently, in vector form, substituting the left hand side of the equation by 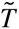, which can be computed from the measured target levels,

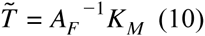

Here *K_M_* is a (1 × *M*)-matrix, 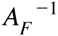 an (*N* × 1)-matrix and 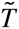 a (*N* × *M*)-matrix. 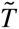 can be viewed as a Kronecker product of the two vectors *K_M_* and 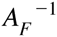, written in a more general form as

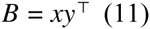

Determining the vectors *x* and *y* becomes a problem known as reverse Kronecker product and has a known solution satisfying

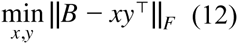

where || · ||*_F_* denotes the Frobenius norm. The solution is obtained from the singular value decomposition (SVD) *B = UΣΥ*^⊤^ as

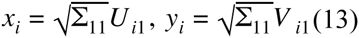

From equation (9) we see that the SVD provides us the solution (*A_F_, K_M_*), up to a scaling factor *a*, 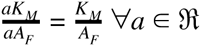, (see **Figure 4:a)**. We note that in principle it is possible to determine the factor *a* which explains best the data, using the total concentration of Ago-miRNA complexes *A* in all cells.

Fitting the vectors *A_F_* and *K_M_* on *in-silico* data, it appeared that the correlation of the fitted *A_F_* with the input value is significantly higher than for *K_M_*. This is explained by the fact that we use the total concentration of the miRNA in the cells to sort the cells and smoothen the mRNA expression. Because *A_F_* is a monotonic strictly increasing continuous function of *A*, the moving average performed along the cells, so the lines of the matrix *T* (i.e. along the *j* index in equation (8)) leads to a reduction of noise in the *A_F_* (cell) dimension but not in the *K_M_* (mRNA) dimension. Therefore, the vector *A_F_* is inferred more precisely compared to *K_M_*. Using the more precisely inferred *A_F_* values and averaging over cells, we can increase the precision of *K_M_* values: relation (10) always holds and after fitting, we use the values of *A_F_* to compute the values *K_M_* by averaging 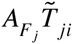 over all cells *j =* 1…*N*

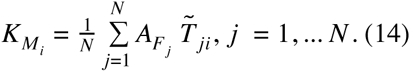

### *In silico* model verification

Stochastic simulations based on equations (1) were used to verify the solution obtained in equation (8) (**Figure 2:a**). Stochastic simulations were performed using StochKit v.2.0.11 [45] with a tau leaping algorithm. For each *in silico* cell, 6 simulations of length 100’000 (arbitrary time units) were carried out to ensure convergence. The first 10’000 steps were considered the ‘burning phase’, and were discarded before the analysis. Means and standard deviations were calculated from the values obtained in the independent simulations.

To test the *K_M_* inference method we constructed an *in silico* data set as follows. We considered a regulatory network of 100 miRNA targets. Each target was characterized by parameters 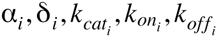, whose average values we assumed to be in the ranges provided by our previous literature survey [11]. For each target we chose a set of parameters from log-normal (or exponential) distributions around the above-mentioned mean values, such that the final set of *in silico* targets resembles the experimental set in the distribution of i) target expression levels and ii) maximal miRNA-induced down-regulation and iii) range of miRNA concentrations where targets are maximally sensitive to the miRNA (compare **Figures 6:e** and **4:j**). The distributions of each individual parameter in this final set of *in silico* targets are shown in **Figure S7**. We considered 300 virtual cells, each with a distinct concentration of free Ago-miRNA complexes, chosen from a uniform distribution on the 1-17 log_2_ range. The expression of all targets as a function of the miRNA abundance in these virtual cells is presented in **Figure 4:b**. Targets are down-regulated around a specific threshold, where the total target population balances the miRNA concentration. The sensitivity of each target, defined by the maximum slope of its decay in function of the miRNA concentration depends on the *K_M_* of the target. The change in target level at saturating miRNA concentrations relative to the situation when the miRNA is not expressed depends only on how much the miRNA accelerates the decay of the target. As observed experimentally (see [11] for a summary), the change is small for most targets (**Figures 5:a, 4:b and 4:c**). To complete our *in silico* data generation we added log-normal noise to the expression levels of the targets that were calculated with the model, to reflect what has been observed in single cell sequencing experiments (see **Figure 4:d**).

The fitting procedure started with the selection of single cells from which to construct the matrix 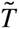 (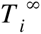 and 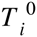 were calculated from a few tens of cells with the lowest and highest concentration of miRNA) because we wanted to focus on cells were the target levels most sensitively responded to the miRNA level in the cell, rather than on cells where the target expression variation was not due to the miRNA. Therefore, we analyzed the derivative of (sum of log_2_) target levels in function of miRNA expression and selected the cells where were the gradient was lowest (**Figure 4:e**). Cells with target expression values very close to 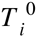 or 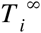 were filtered out to avoid instabilities caused by division by small numbers (see Equation 9). Next, we sorted the cells according to the miRNA expression and applied smoothing procedure to ensure that at intermediate miRNA expression range, the *T_ij_* level of targets *i* in cells *j* was strictly in the range 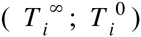 (see **Figure 4:f**). We started by replacing the target expression level in one cell with the median among the cell and its five closest neighbors (in terms of miRNA expression). We then computed the running mean starting with a window size of ten, and discarded iteratively the strongest outliers until the mean value T_ij_ within each window was within the 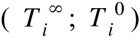 range. For the windows were this procedure did not leave any points, we increased the window size locally and repeated the pruning procedure until all the T_ij_ values were within the 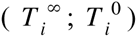 range. To ensure the stability of the SVD we adjusted the boundary of the T intervals computed from the data by a small safety margin c (i.e. 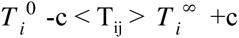).

We assessed the accuracy of the fitting procedure by comparing the inferred *K_M_* parameters and miRNA concentrations with those that were used in the model that generated the *in silico* data. In spite of very high noise (**Figure 4:d,f**) there is a good correlation between the fitted and input values of the parameters, as shown in **Figure 4:g-h**. This is probably due to the fact that the 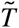 matrix has ~ 10^4^ elements and each parameter comes as part of an optimal solution for this large matrix; in principle, if the data were extremely accurate, only two cells would be sufficient to infer *K_M_* constants (aside from two control cells, in which the miRNA is not expressed and expressed at saturating concentrations, respectively). However, even with the *in silico* data the correlation of the parameters inferred from two instantiations of the model in which the target expression levels differed due to independent application of “experimental” noise is limited (**Figure 4:i)**. This test provides a realistic estimate of the correlation that we expect between parameter values inferred from two experimental data sets. We also observed that the range of inferred *K_M_*s is narrower than the range of input *K_M_*s. This suggests that the real range of variation of *K_M_* values is probably larger than we can infer with our method (**Figure 4:k**). The average *K_M_* is preserved, and the difference in the response characteristic of 10 targets with the lowest and 10 with highest *K_M_*s is clear for both the original and the inferred sets (**Figure 4:j**).

Having validated our inference procedure on *in silico* data, we applied it to the experimental data.

### Cell culture

For this study we used a Human Epithelial Kidney (HEK) 293 cell line with inducible expression of hsa-miR-199a (i199) and a derivative of this cell line (i199-KTN1) in which a Renilla Luciferase reporter gene with the 3’UTR of the kinectin 1 gene (KTN1) was inserted in the genome. These cell lines have been introduced before [6]. Cells were grown in DMEM media with 10% FCS supplemented with Pen-Strep and Hygromycin for plasmid integrity. For all the experiments, unless otherwise mentioned, cells were stimulated with doxycycline, at the corresponding doses, for 8 days. During this period, fresh medium with doxycycline was provided every 24 hours.

### Single cell mRNA-sequencing

#### Cell capture, imaging and cDNA synthesis

Cells were detached with Accutase^®^ reagent (Gibco, Life Technologies ™). The cell number was determined with the Countess™ Automated Cell Counter (invitrogen ™) following manufacturer’s instructions. Cells that were induced with different doxycycline conditions were pooled together in equal proportions, to a concentration of 3 million cells per ml. The mixed cell suspension was diluted further to 300’000 cells per ml. 5 μl of the last suspension was used to load the AutoPrep IFC microfluidic chip (C1™ Single-Cell Auto Prep IFC for mRNA Seq (10-17 μm) PN-100-5760, Fluidigm Corporation) according to the manufacturer’s protocol. After loading, each capture site was checked by microscopy for capture efficiency. The chip was further processed on the Fluidigm C1 instrument to obtain the corresponding cDNA, using the standard C1™ mRNA-Seq built-in program and the SMARTer^®^ Ultra™ Low RNA Kit for the Fluidigm C1™ System (Clontech, Inc.) reagents. Briefly, cells undergo lysis, reverse transcription with template switch and pre-amplification of the double stranded cDNA by means of 21 PCR cycles. The pre-amplified cDNA is then transferred to a 96 well plate, for further processing.

#### Library preparation

Pre-amplified cDNA was simultaneously fragmented, amplified and barcoded using Nextera XT DNA Library Preparation Kit (Illumina inc.) following manufacturer’s protocol, adapted by Fluidigm^®^ for low input material. Briefly, the cDNA concentration was measured with Picogreen^®^ (Life Technologies, ™) and the cDNA was diluted to a concentration of 0.1-0.3ng/μl. Diluted cDNA underwent tagmentation with Amplification Tagment Mix reagent, which adds specific adaptors while fragmenting the cDNA. Tagmentation was done at 55°C for 10 min. After tagmentation enzymes were neutralized with neutralizing buffer, the fragmented cDNA was amplified by 12 cycles of PCR. Two different barcodes were added to each sample to achieve 96 different barcode combinations. The libraries of each cell were combined together and cleaned up from the unused PCR primers by double Agencourt AMPure XP (Beckman Coulter, Inc.) bead purification, with a 0.9x ratio.

#### Library sequencing

The libraries were sequenced in the Genomics Facility Basel, on Illumina HiSeq 2000, HiSeq 2500 or NextSeq instruments using Nextera XT compatible primers. Reads of 50-76 nt were generated along with dual index reads corresponding to the cell-specific barcodes. The data has been deposited to the Sequence read archive (www.ncbi.nlm.nih.gov/sra/) under the accession number SRP067502.

### Cell population mRNA-seq

#### Total RNA isolation

Total RNA was extracted with TRI Reagent^®^ (Sigma-aldrich) following manufacturer’s instructions. Briefly, cells were detached from the plate by 5 min incubation with Trypsin-EDTA solution (T3924 SIGMA), conditioned media was added and whenever necessary, cells were counted with a Countess™ cell counter (Thermo Fisher Scientific). A defined number of cells were pelleted and either snap frozen for future use or resuspended right away. After rapid thawing, 1 ml of TRI Reagent was added to lyse the cells and extract total RNA. Total RNA was resuspended with DEPC-MQ-H2O. Samples were always kept on ice or at −80°C.

#### mRNA purification

To select the Poly(A)^+^ RNA, a double purification with Dynabeads^®^ Oligo (dT)25 (Dynabeads^®^ mRNA DIRECT™ Kit, Ambion™) was performed, using the manufacturer’s manual and recommendations. Since the starting material was purified total-RNA, only buffer B was used for the washing steps.

#### Library preparation

Purified mRNA was fractionated with Alkaline Hydrolysis Buffer at 95°C for 5 min. Fractionated mRNA was selected with RNeasy MinElute Cleanup Kit (Qiagen, Inc.). Purified mRNA fragments were dephosphorylated with FastAP (Life Technologies, Inc.) and 5’-phosphorylated with PNK (Life Technologies, Inc.) following manufacturer’s instructions for optimal conditions of the enzymatic reaction. After another round of RNeasy MinElute Cleanup Kit (Qiagen, Inc.), a pre-adenylated DNA adapter (5’-TGGAATTCTCGGGTGCCAAGG-3’) was ligated to the 3’end of the mRNA fragments at 4°C overnight using the T4 RNA ligase 2, truncated K227Q (New England Biolabs, Inc.), in 1x T4 RNA ligase buffer (no ATP) and 15% DMSO. The next day, after another round of RNeasy MinElute Cleanup Kit (Qiagen, Inc.), an RNA adapter (5’-GUUCAGAGUUCUACAGUCCGACGAUC-3’) was ligated to the 5’end of the RNA fragments at 4°C overnight using the T4 RNA ligase 1 (Life Technologies, Inc.), in 1x T4 RNA ligase buffer (1mM ATP) and 15% DMSO. Next day, after another round of RNeasy MinElute Cleanup Kit (Qiagen, Inc.), Reverse Transcription was performed using Superscript IV (invitrogen, Inc.) and RTP primer (5’-CCTTGGCACCCGAGAATTCCA-3’), following manufacturer’s instructions. cDNA was then amplified by 12 cycles of PCR using NEBNext^®^ High-Fidelity 2X PCR Master Mix (NEB, Inc.), and Illumina TruSeq^®^ Small RNA PCR compatible primers.

#### Library sequencing

The library was sequenced in the Genomics Facility Basel, on Illumina HiSeq 2000 or HiSeq 2500 instruments using Truseq compatible primers. Reads of 50 nt were generated along with 8nt index reads corresponding to the sample-specific barcode.

### Read mapping and data preprocessing

Reads from single cell and cell population mRNA-Seq were mapped to the transcriptome (Ensembl, GRCh38.rel79) with kallisto v0.42.1 [46]. The sequence of the eGFP mRNA was added to the transcriptome before read mapping. Since in single cell sequencing the fragment length varies widely and is not normally distributed, we set the fragment length to a small number (50/75, equal to the read length) for all the sets. The net outcome of this choice should be effectively turning off effective length correction in the estimation of read coverages. We have also used another software tool, sailfish [47] (version 8.0, without taking into account the transcript length by using option *noEffectiveLengthCorrection*), to quantify transcript expression and we obtained similar results. Finally, we also explored the change of target prediction method [48]. All of these variations lead to similar results (see fitting of *K_M_*s obtained with a different set of methods in **Figure S10**).

To obtain obtain gene-level expression values, we summed the estimated transcript expression levels (transcripts per million, TPM) provided by kallisto. For each gene the values that were more than 3.5 (i199-KTN1 line) standard deviations away from the mean over the entire set of cells were set to NaN. We also excluded from the analysis cells whose expression values had a median correlation with the values of other cells below 0.64 (5-10% of the cells, see **Figure S9**) and whose total gene expression level was less than ~500’000 TPM. In all the analysis genes with very low expression (mean TPM <15) were not used. For the validations presented in **Figure 1**, control cells were considered those with log_2_ GFP expression lower than 8 and and induced cells were those with log_2_ GFP expression higher than 12.

#### Targets selection

In both cell lines we studied (if not specified otherwise) the response of the 200 highest probability targets predicted by MIRZA-G-C [31] for each of the two distinct miRNAs, that were down-regulated at least 10% at the maximum miRNA concentration 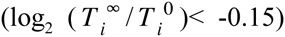. Additionally we required the targets to be downregulated in the induced cells compared to the control (as specified in the paragraph above, with the log_2_ ratio of the median values < −0.15), to avoid targets with high fluctuations (in function of GFP expression) of the expression levels.

### mRNA and miRNA qPCR

Remnant cells from the C1™ single cell mRNA-seq experiments, as well as additional cells derived from the same experiments, were used for the qPCR assays. After counting the cells, total RNA was extracted with TRI Reagent^®^ (Sigma-aldrich) following manufacturer’s instructions. cDNA of the targets of interest was generated using superscript III (Invitrogen ™) following manufacturer’s protocol. For miRNA assays, reverse transcription and PCR of either non-induced or Dox induced cells were performed following the TaqMan^®^ Small RNA Assays quick reference protocol (Life Technologies™) with 100ng of total RNA. For estimation of relative miRNA quantities, hsa-miR-16 levels were used as an invariant control. For reverse transcription of GFP mRNA, the following linear DNA primer was used: EGFP_R RT taqman, 5’-TGTCGCCCTCGAACTTCAC-3’. To generate a cDNA copy of hsa-miR199a-5p a stem-loop primer system from Life technologies ™ was used (Assay ID-000498). All qPCR were performed and read in StepOnePlus™ Real-Time PCR Systems (Life Technologies™). To obtain absolute quantification data, standard curves for GFP and hsa-miR-199a-5p were also included. GFP mRNA was generated by in-vitro transcription with pcDNA3-eGFP linearized vector and RiboMAX™ Large Scale RNA Production System - T7 (Promega, Co.) following manufacturer’s instructions. Molarity was estimated taking into account mass concentration (Qubit^®^ RNA HS assay kit - Life Technologies ™), average length (Agilent RNA 6000 Pico Kit - Agilent Technologies, Inc) and fragment sequence, with the following formula: molarity = mass/(length * mass RNA base). The hsa-miR-199a-5p miRNA (5’-CCCAGUGUUCAGACUACCUGUUC-3’) was ordered from Microsynth AG, and the molarity was calculated the same way. Absolute molecule numbers were obtained utilizing the StepOne™ Software (Life Technologies™).

#### Weighted sum of qPCR measurements

For a given doxycycline concentration *i*, the the weighted mean expression level 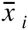 of the analyzed miRNA targets *j* was calculated using the formula below

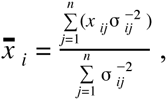

where σ*_ij_*, is a standard deviation calculated from the biological replicates of a measurement.

## Supplementary plots

**Figure S1.**
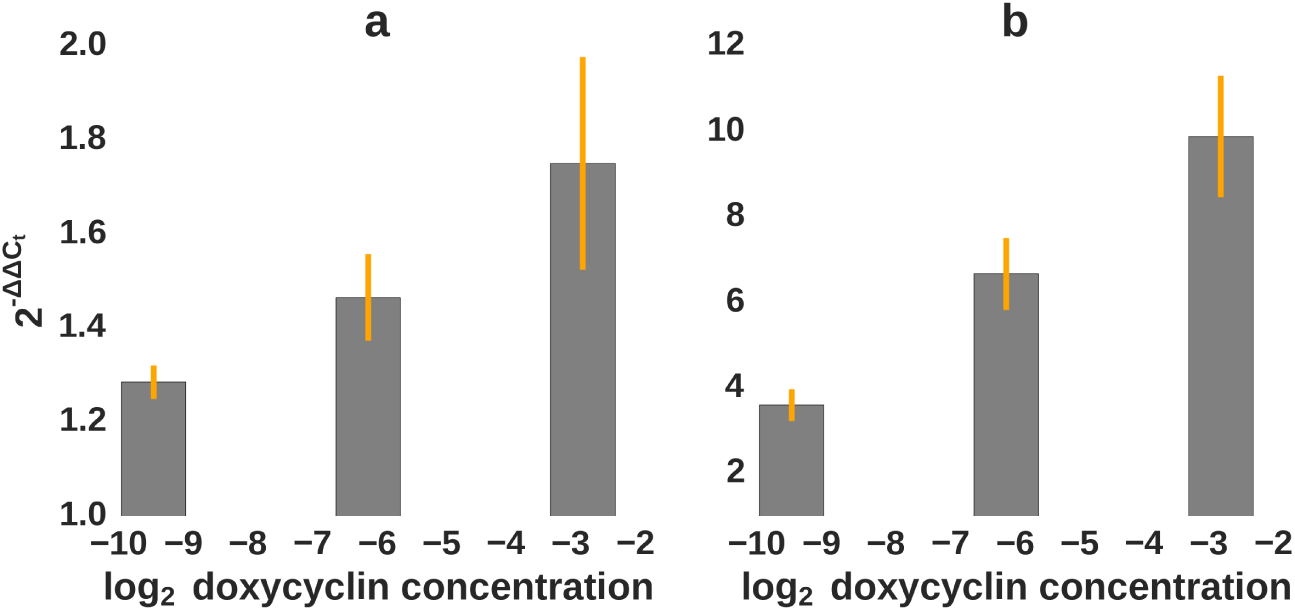
Both hsa-miR-199a-5p and hsa-miR-199a-3p are processed from the hsa-miR-199a precursor. miRNA induction measured with quantitative PCR. **a-b)** Relative hsa-miR-199-3p (a) and hsa-miR-199-5p (b) miRNA levels in doxycyclin-induced cells compared to the non-induced cells. The C_t_ values obtained for each set were normalized to the levels of hsa-miR-16 and to the values from non-induced cells.

**Figure S2.**
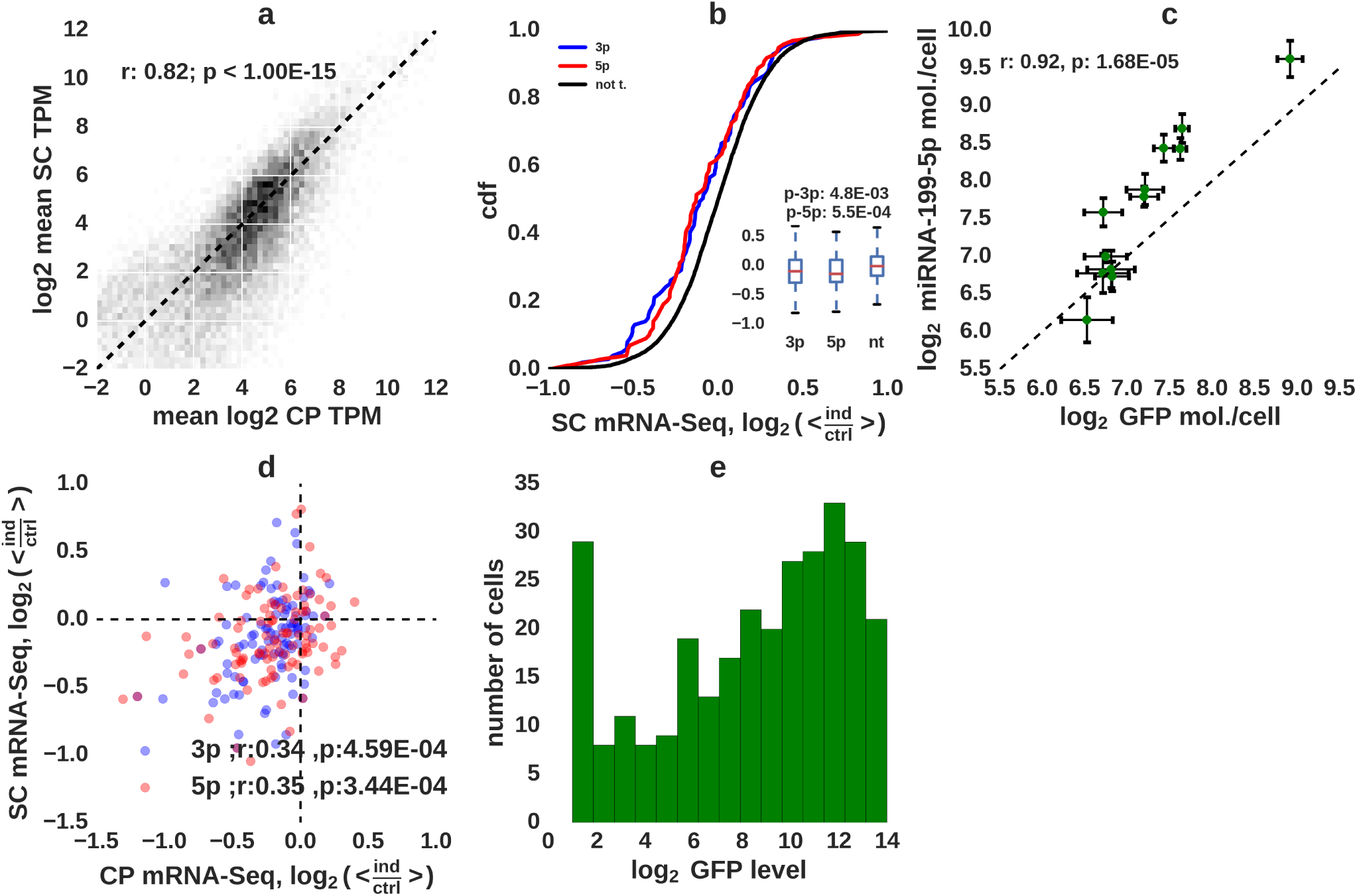
Characterization of miRNA activity in single i199 HEK cells. **a)** Correlation of individual mRNA expression levels measured with SC (~170 cells that did not express the miRNAs) and CP (6 replicates of non-induced cell populations). **b)** Downregulation of top 100 targets of each miRNA as inferred with SC mRNA-seq (normalized fold changes, blue: targets of hsa-miR-199a-3p, red: targets of hsa-miR-199a-5p, black: other transcripts). **c)** Correlation of hsa-miR-199a-5p and GFP mRNA expression measured in cell populations by quantitative PCR. **d)** Correlation of downregulation in SC and CP mRNA-Seq for 100 top targets. **e)** GFP mRNA expression distribution in single cells.

**Figure S3.**
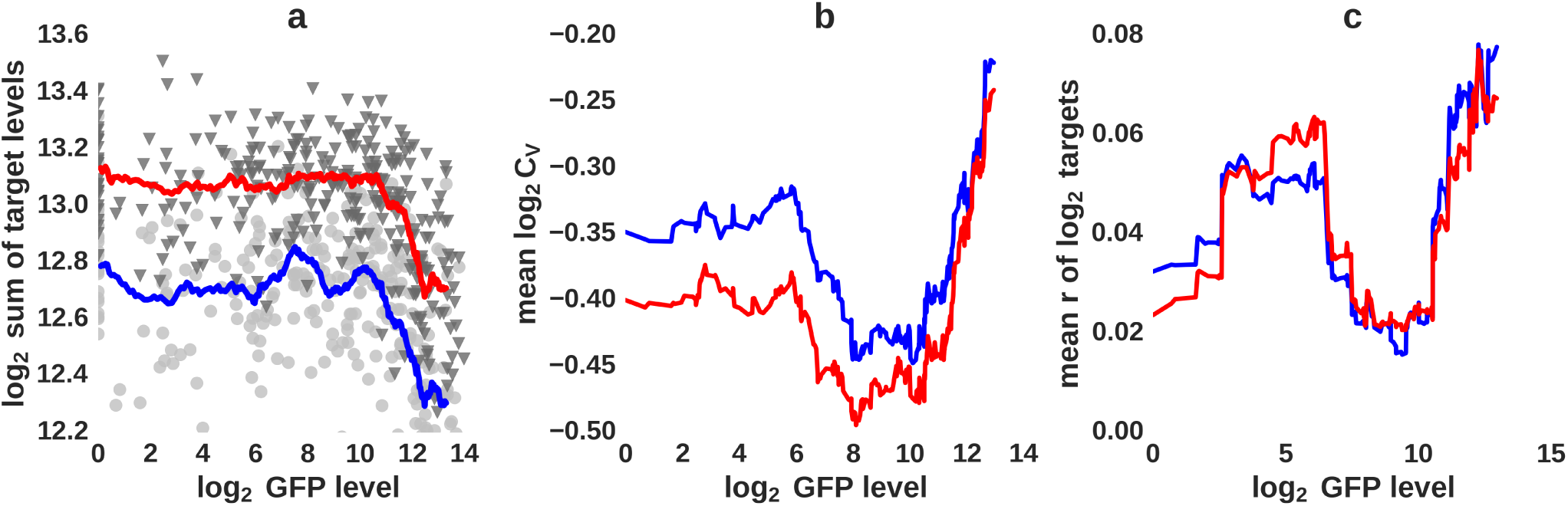
Inferring the effect of miRNAs on mRNA targets from single cell mRNA-seq. Target selection is described in the main text, shown are the results obtained on i199 cells. **a)** Total target expression (log_2_sum of TPMs of hsa-miR-199a-3p and hsa-miR-199a-5p predicted targets, see Methods for target selection) in i199-KTN1 cells in function of log_2_ GFP expression. The colored lines show running mean with window of 30 cells, the grey dots and black triangles show the expression of the 3p and 5p targets, respectively, in individual cells **b)** Mean C_V_ and **c)** mean Pearson pair-wise correlation coefficients for miRNA targets in function of GFP expression. For each GFP level averages were calculated from the fifty cells with GFP expression closest to the reference level.

**Figure S4.**
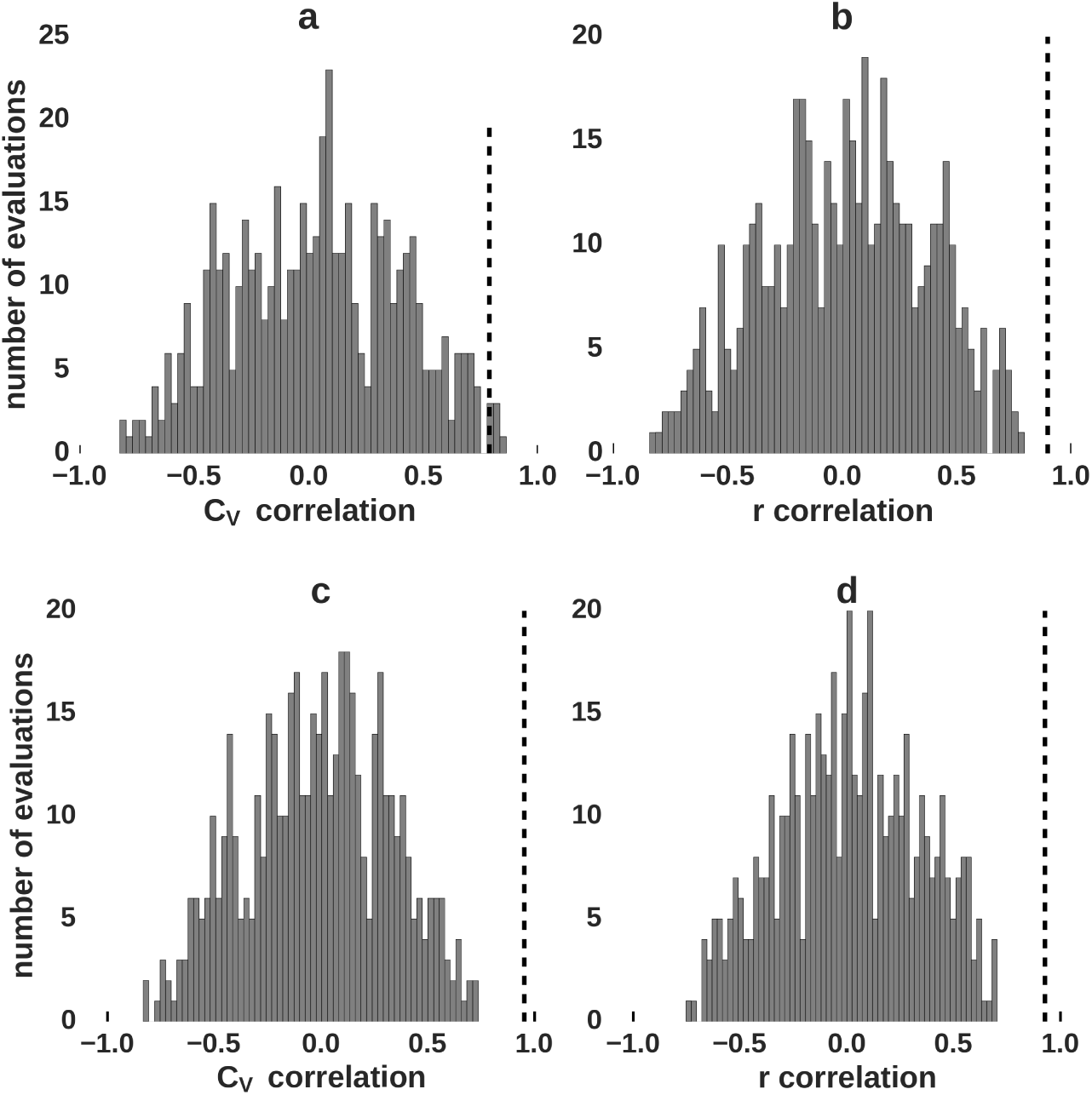
Evaluation of the significance of noise and target correlation signatures of the two miRNAs in single cells. The cross-correlation between the *C_V_* and *r* profiles of targets of the 3p and 5p miRNAs in function of GFP/miRNA expression were calculated for the experimental data set (shown in **Figure 3:b,c)**. These values are indicated by the vertical dashed line. The cells were then shuffled to randomize their miRNA expression and the procedure was repeated. Shown are Spearman correlations between *C_V_* (**a,c**) and r profiles (**b,d**) in function of (randomized) GFP/miRNA levels for the two sets of targets. The histograms show results for 500 independent evaluations. **a,b)** show the results for the i199-KTN1 line, **c,d)** for the i199 line.

**Figure S5.**
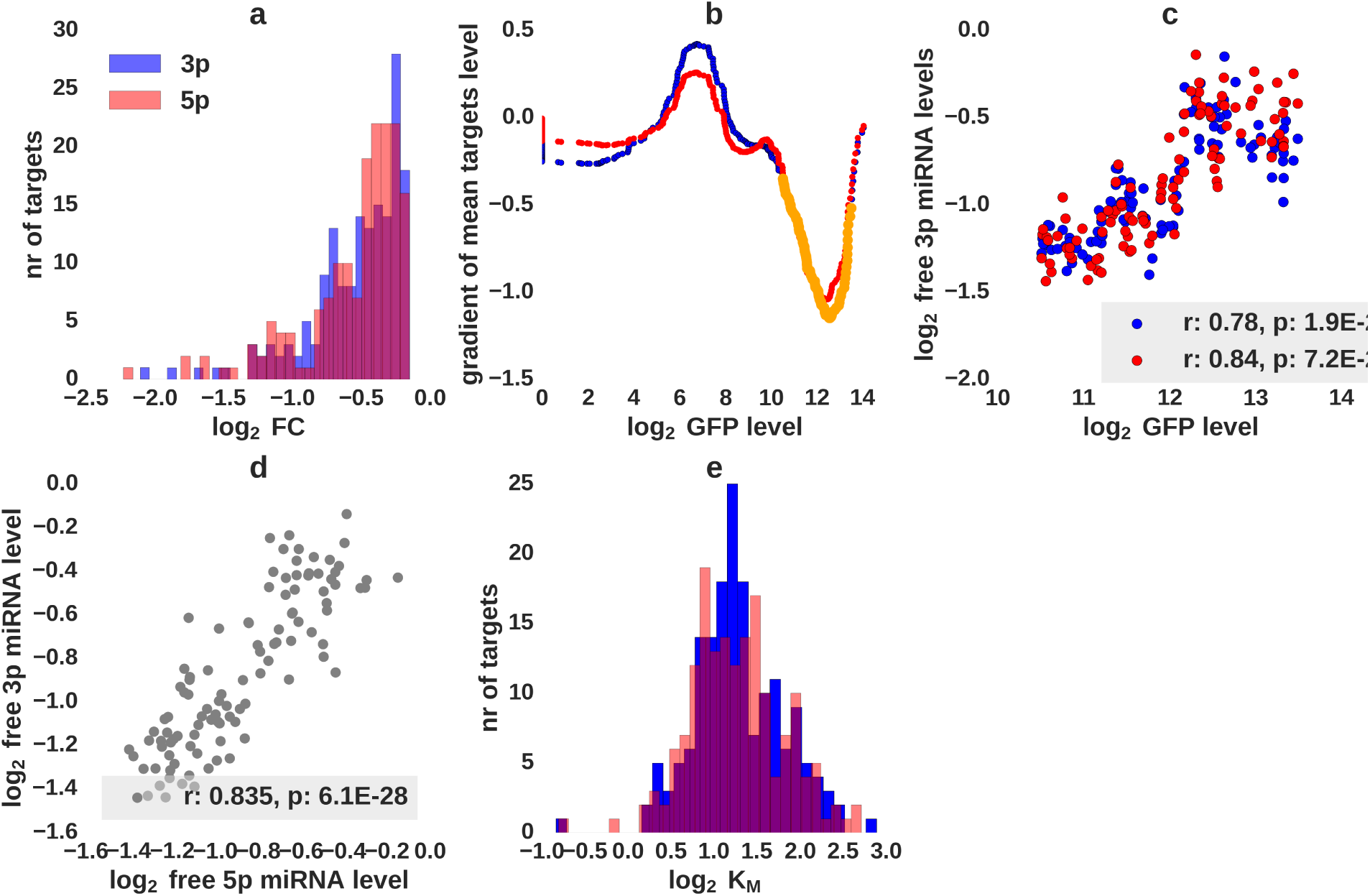
Fitting *K_M_* constants from SC mRNA-seq expression measurements. Results for the i199 cell line. **a)** Selection of targets. Targets with log_2_ fold change < −0.15 at maximal miRNA concentration were considered. **b)** Selection of cells for the inference of *K_M_*s. Target levels in cells with log_2_ GFP expression in the range of 10-13 (orange line) were used to construct the *T* matrix. **c)** Spearman correlation of the inferred levels of free miRNAs 5p and 3p and the measured GFP mRNA. **d)** Pearson correlation of the inferred levels of free 5p and 3p miRNAs. **e)** Distribution of *K_M_* values.

**Figure S6.**
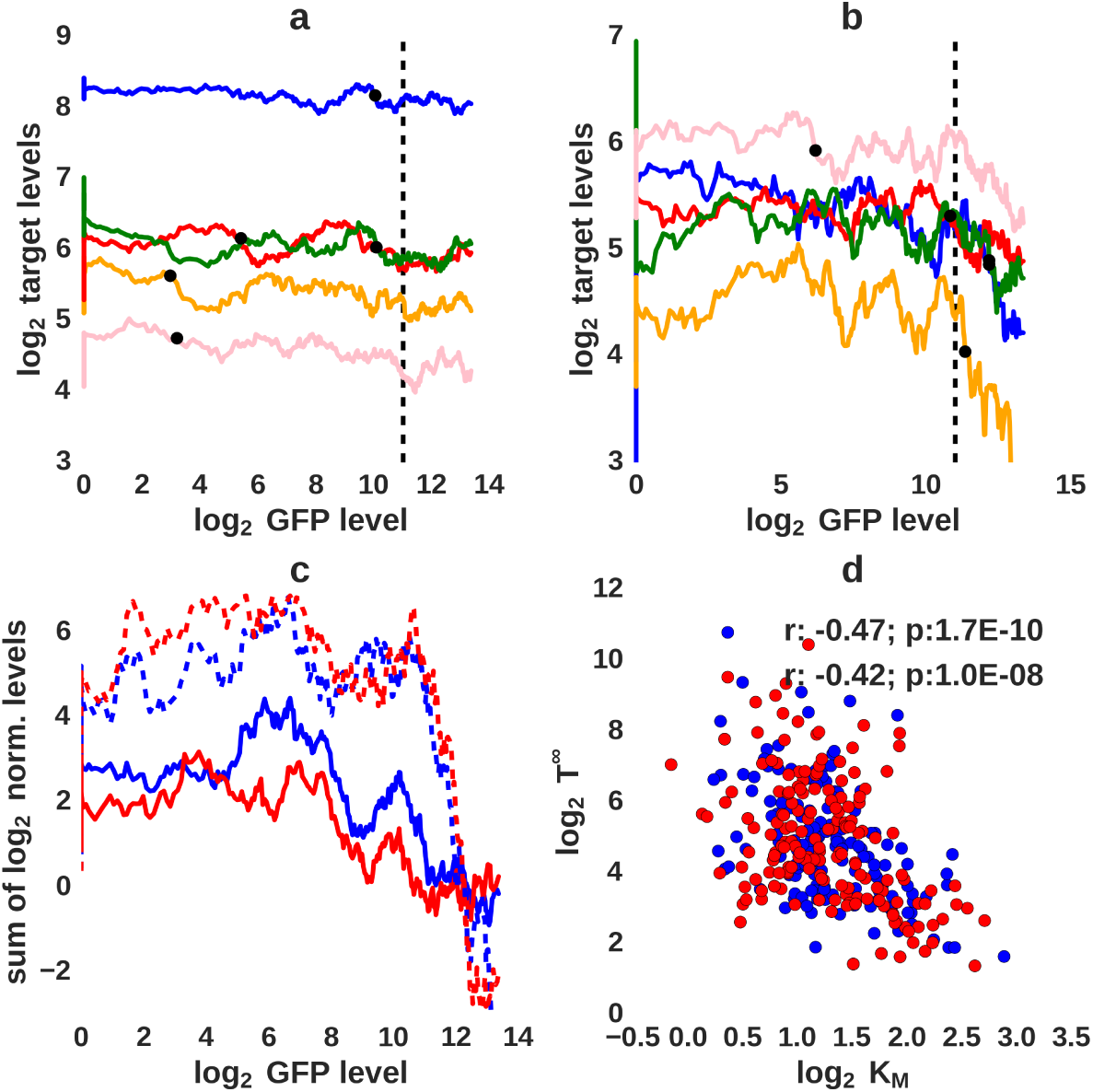
Validation of *K_M_*s fitted based on the single cell data. **a)** Single cell level expression of five low *K_M_* miRNA targets in function of the GFP mRNA. Shown are the running means (window size of 30 cells, black dots show position of the minimum value of the gradient). **b)** As in **a**, but for five high *K_M_* miRNA targets. **c)** Sum of log_2_ running means (normalized before summation by the corresponding 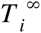 values) for ten targets with the lowest (solid line) and highest (dashed line) inferred *K_M_* values. Red and blue correspond to targets of the hsa-miR-199a-5p and hsa-miR-199a-3p miRNAs, respectively. **d)** Scatter plot of *K_M_* and 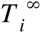 for individual targets of the 5p (red) and 3p (blue) miRNAs. r is the Spearman correlation coefficient.

**Figure S7.**
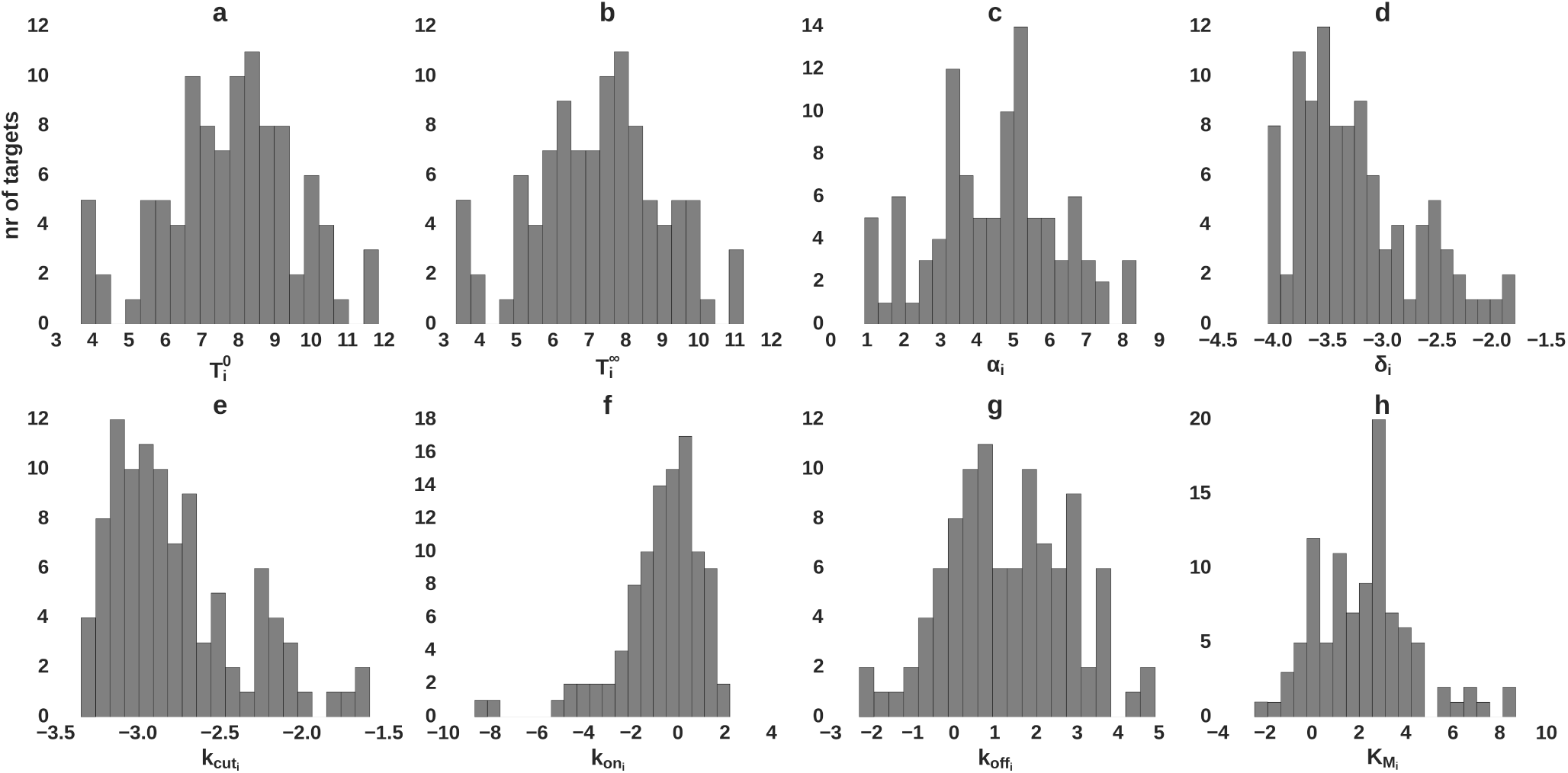
Distribution of parameters for *in silico* targets. Log_2_ values are shown. See section *In silico* model verification for additional explanation.

**Figure S8.**
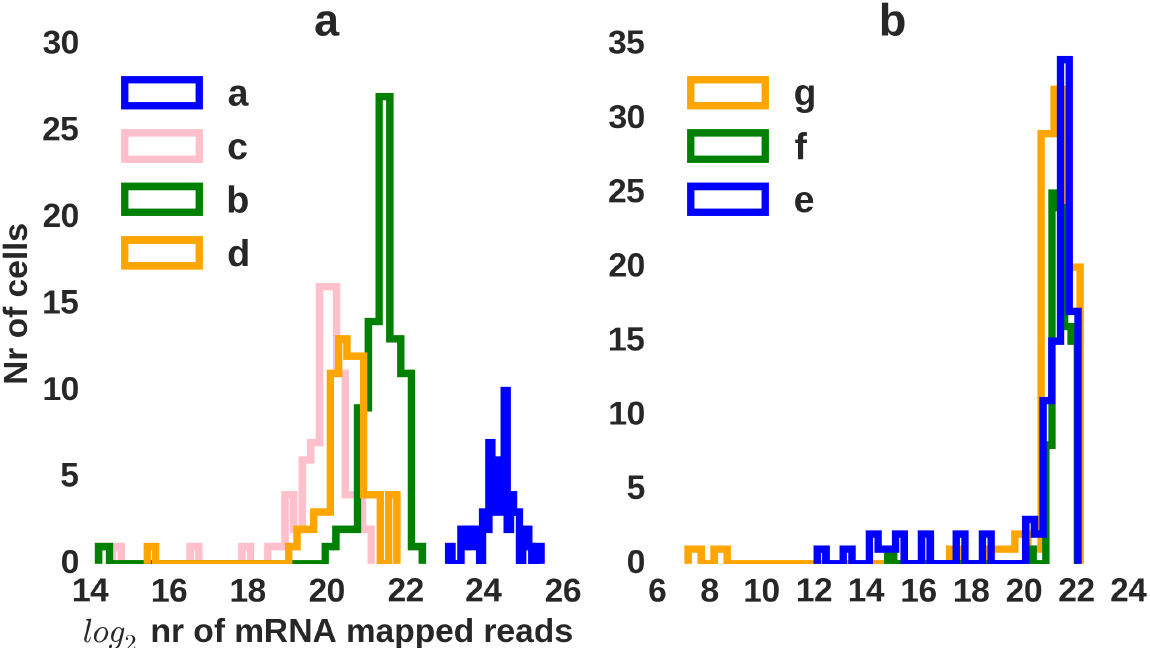
Histograms of the number of reads obtained from individual cells, in the different sequencing experiments (a-e) that were mapped to mRNAs. **a)** Experiments with the i199 cell line, **b)** Experiments with the KTN1-199 cell line. Mapping was performed to the transcriptome (including the GFP transcript) and genome, and only the lowest-error mappings were used (see [49]).

**Figure S9.**
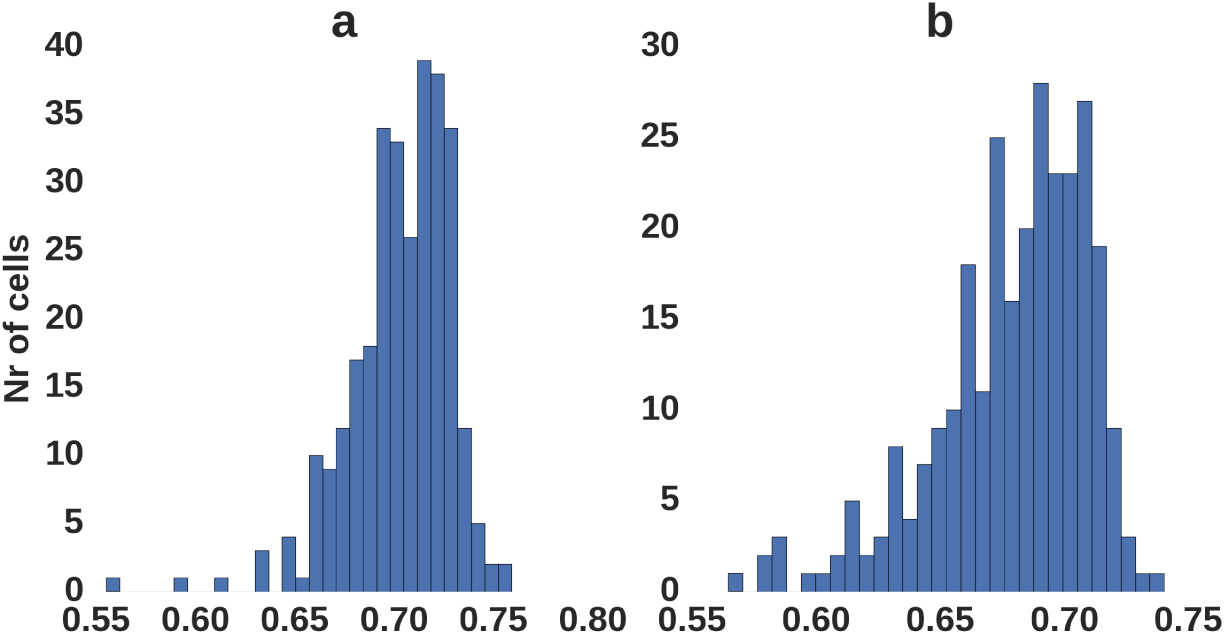
Correlation of gene expression levels inferred from individual cells. Histogram of the median Pearson correlation of log_2_ gene expression levels of a cell with all the other cells. **a)** Data for the i199 line, **b)** Data for the KTN1-199 line.

**Figure S10.**
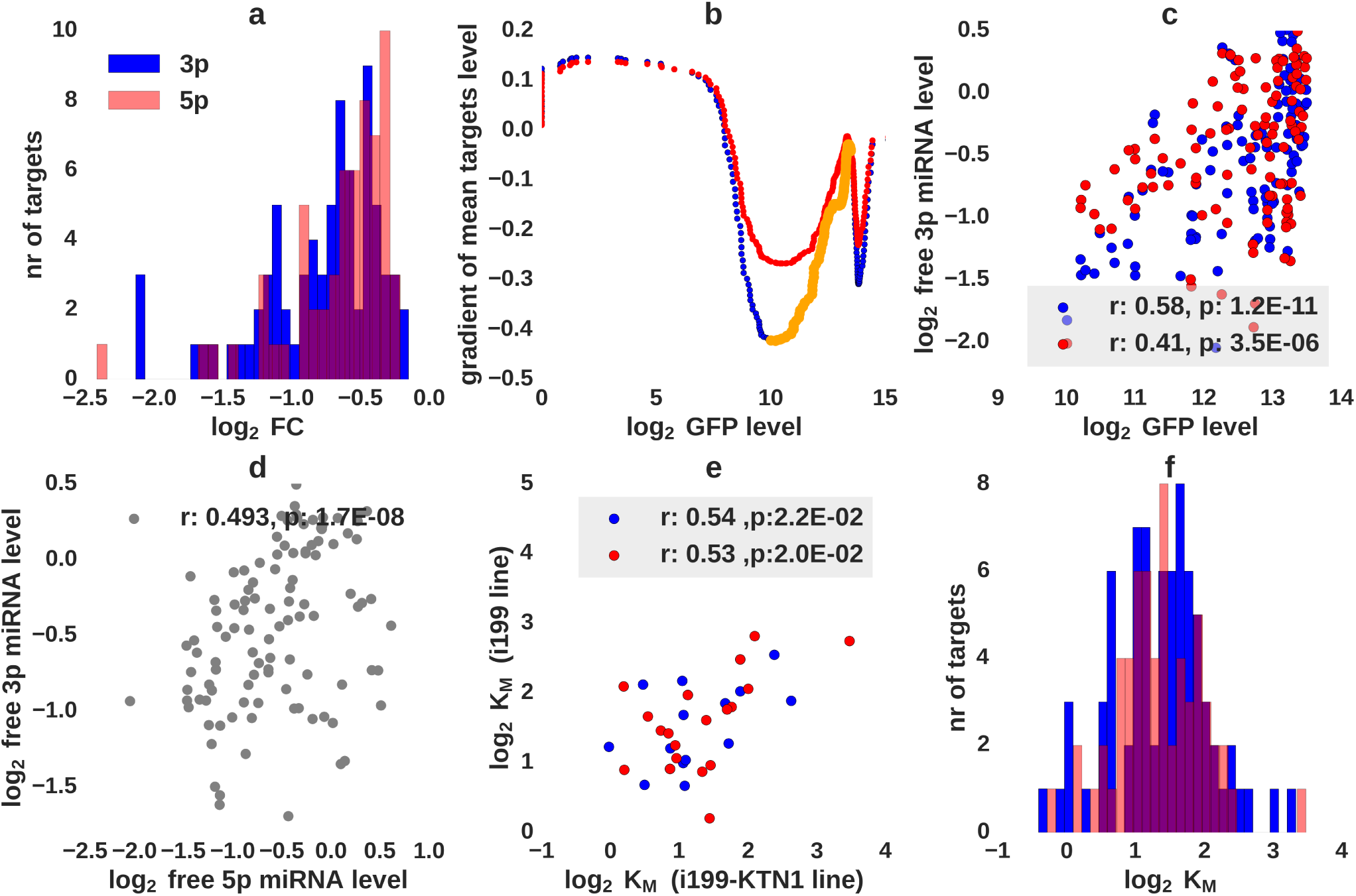
Fitting *K_M_* constants from SC mRNA-seq expression measurements is insensitive to the expression estimation method and the target prediction tool. We performed the same calculations as in Figure 5, but with expression levels obtained from sailfish [47] and targets predicted with Target Scan v.7.0 [48]. Additionally *K_M_* fitting was performed on a transcript level (without summation of transcripts of the same gene). Small modifications of the fitting procedure parameters (i.e. transcripts with expression > 10 TPM rather than 15 were considered, since genes have higher expression) were included to improve the fit results. **a)** Selection of targets. Targets with log_2_ fold change < −0.15 at maximal miRNA concentration were considered. **b)** Selection of cells for the inference of *K_M_*s. Target levels in cells with log_2_ GFP expression in the range of 10-13 (orange line) were used to construct the 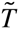 matrix. **c)** Spearman correlation of the inferred levels of free miRNAs 5p and 3p and the measured GFP mRNA. **d)** Pearson correlation of the inferred levels of free 5p and 3p miRNAs. **e)** Pearson correlation of *K_M_* values inferred for individual targets from the i199 and i199-KTN1 cell lines. **f)** Distribution of *K_M_* values.

**Table S1.** Doxycycline concentration that were used in different experiments, grouped according to the C1 single cell chip that was used to capture the cells and prepare the libraries.

| Single cell chip (up to 96 cells) | Doxycycline concentration ($\mu\text{g/ml}$) |
| --- | --- |
| i199 (a) | 1.0, 0.0 |
| i199 (b) | 0.003, 0.0003, 0.0001, 0.00003 |
| i199 (c) | 0.001, 0.0003, 0.0001, 0.00001 |
| i199 (d) | 0.01, 0.001, 0.0003, 0.0001, 0.0 |
| i199-KTN1 (e) | 1.0, 0.1, 0.01, 0.001 |
| i199-KTN1 (f) | 0.3, 0.03, 0.003, 0.0003, 0.0 |
| i199-KTN1 (g) | 0.001, 0.0003, 0.0001, 0.00003 |

